# Cyclin N-Terminal Domain-Containing 1 (CNTD1) coordinates meiotic crossover formation with cell cycle progression in a cyclin-independent manner

**DOI:** 10.1101/750596

**Authors:** Stephen Gray, Emerson R. Santiago, Joshua S. Chappie, Paula E. Cohen

**Affiliations:** Department of Biomedical Sciences and Center for Reproductive Genomics, Cornell University, Ithaca, NY, 14853, United States of America; Department of Molecular Medicine, Cornell University, Ithaca, NY, 14853, United States of America

**Keywords:** Meiosis, Crossing over, CNTD1, Cullin RING ubiquitin ligase, SCF, RFC, PCNA, CDC34, WEE1, CDK1, CDK2, MutLγ, Ubiquitylation

## Abstract

During meiotic prophase I, programmed DNA double-strand breaks repair as non-crossover or crossover events, the latter predominantly occurring via the Class I crossover pathway and requiring the cyclin family member CNTD1. Using an epitope-tagged *Cntd1* allele, we show that mouse CNTD1 exists *in vivo* as a short isoform that lacks the predicted N-terminal cyclin domain and does not bind cyclin-dependent kinases. Instead, we find that CNTD1 associates with Replication Factor C to drive crossover formation and the Skp1-Cullin1-F-Box complex to regulate ubiquitination and subsequent degradation of the WEE1 kinase, thereby indirectly modulating cell cycle progression. We propose that these interactions enable CNTD1 to orchestrate the steps of prophase I and coordinate crossover formation with cellular division.

## Introduction

Meiosis is a specialized cell division consisting of one round of DNA replication followed by two rounds of chromosome segregation. During meiosis I, homologous chromosomes must pair, undergo physical tethering (synapsis), and form crossovers to enable accurate segregation at the first meiotic division (MI). During the second meiotic division (MII), the paired sister chromatids segregate, resulting in the formation of haploid gametes^1^.

Crossing over is achieved through the process of meiotic recombination, which is dependent on, and required for, accurate synapsis. Synapsis is achieved by the formation of the synaptonemal complex (SC), whose assembly and status defines the five sub-stages of prophase I: leptonema, zygonema, pachynema, diplonema, and diakinesis. Meiotic recombination is initiated by the formation of programmed DNA double-strand breaks (DSBs) throughout the genome in early leptonema^2–7^. In the mouse, 200-300 DSBs are generated and processed through common intermediate repair steps to yield either non-crossover or crossover events^1^. While the majority of DSBs (∼90% in the mouse) repair as non-crossovers in zygonema and early pachynema, the crossovers that form during pachynema establish the inter-homolog tethers that allow for their correct segregation at MI^8^. Thus, the correct timing, frequency, and distribution of crossovers ensures that at least one crossover per pair is generated (the obligate crossover)^9^, that crossovers are maintained at the expense of non-crossovers (crossover homeostasis)^10, 11^, and that the formation of one crossover prevents the formation of nearby crossovers (interference)^12^. How these rules are enforced within the realm of DSB repair and shape crossover/non-crossover decisions remains unclear.

In the mouse, crossovers can form by at least two mechanisms. The class I (ZMM) pathway is responsible for the majority of crossovers and utilizes the DNA mismatch repair endonuclease MutLγ, a heterodimer of MLH1 and MLH3^13, 14^. MLH1 and MLH3 co-localize along chromosome cores at a frequency of approximately 23 foci at pachynema^15^, a number that reflects approximately 90-95% of the final crossover count in male meiosis^14, 16^. The distribution of MutLγ foci during pachynema in mouse spermatocytes occurs in an interference-dependent manner, at the same time satisfying the requirement for the obligate crossover ^17^. Loss of either component of MutLγ results in infertility due to the loss of most crossovers, leading to meiosis I non-disjunction^13, 14, 18^. The remaining crossovers formed are thought to be generated by the class II MUS81/EME1-dependent pathway. These class II crossovers are interference independent, but cannot account for all the class I-independent crossovers since mutation of both *Mlh3* and *Mus81* leads to the persistence of between 1-3 chiasmata^19^.

One of the major questions in mammalian meiosis concerns how crossovers are selected from the initial pool of 200-300 DSB repair intermediates. Initially, a subset (∼150) of these repair intermediates accrue the MutSγ heterodimer of MSH4 and MSH5^20, 21^, an event termed crossover licensing. Of these, only 23-26 MutSγ sites subsequently become loaded with MutLγ to form class I crossovers while the remaining sites are repaired either through the class II MUS81/EME1 crossover pathway or via the formation of non-crossovers. The mechanism by which MutSγ becomes further selected by accrual of MutLγ has been termed crossover designation, leading to the idea that crossover homeostasis is imposed sequentially by the association of these pro-crossover MutS/MutL proteins^10, 22, 23^.

Recent studies have revealed a number of regulatory molecules that aid in crossover designation and that are essential for class I crossovers, including Crossover site-associated-1 (COSA-1) in *Caernohabditis elegans*^24^ and its mammalian ortholog Cyclin N-terminal domain-containing-1 (CNTD1)^25^. Loss of COSA-1 in worms results in a failure to accumulate MSH-5 at DSB repair intermediates and consequent loss of all crossovers^24^. Loss of CNTD1 results in similar meiotic failure characterized by persistently elevated early crossover factors through pachynema and a failure to load crossover designation factors like MutLγ, the crossover site-associated cyclin-dependent kinase-2 (CDK2), and the putative ubiquitin E3 ligase HEI10 onto DSB repair sites^25^.

CNTD1 and COSA-1 are both distant members of the cyclin family^24^, suggesting that they bind one or more CDKs. To further investigate the function of CNTD1 in mouse meiosis, we generated a null allele of *Cntd1* (*Cntd1^-/-^*) to confirm that loss of *Cntd1* phenocopies our original gene-trap allele (*Cntd1^GT^).* We also generated a C-terminal epitope tag of *Cntd1* (*Cntd1^FH^)* to offset the lack of an effective anti-CNTD1 antibody. Surprisingly, analysis of this tagged allele reveals that the predominant form of CNTD1 in the mouse testis lacks a critical region of the N terminal cyclin domain that is required for CDK interaction. Sequence comparison across an array of species reveals variable splicing and conservation of the CNTD1 N-terminus between divergent orthologs. We demonstrate that CNTD1 forms discrete foci along pachytene chromosomes, co-localizing with MutLγ. We also define the expression profile of CNTD1 relative to other meiotic proteins during prophase I and perform stage-specific mass-spectrometry to identify CNTD1-interacting proteins. Importantly, the mass-spectrometry did not reveal any CNTD1-CDK interaction, in line with our yeast-two hybrid data. Instead we find CNTD1 interactions with components of the Replication factor C (RFC) complex, which functions with PCNA in somatic cells to activate the endonuclease activity of MLH1-PMS1 in DNA mismatch repair^26^. In addition, CNTD1 interacts with components of the major Cullin1 ubiquitin ligase complex (SCF), consisting of SKP1, Cullin 1, a meiosis-specific F-Box protein (FBXW9), and other associated proteins, whose targets include the WEE1 kinase that prevents cell cycle progression into metaphase. Taken together, our studies show that CNTD1 functions not as a cyclin-CDK constituent, but rather as a master regulator that coordinates the activities of two distinct pathways to ensure progression of cell cycle only occurs when the appropriate number of crossover events has been achieved.

## Results

### Epitope tagging of CNTD1 to create a *Cntd1^FLAG-HA^*allele reveals a short form CNTD1

Using CRISPR/Cas9, we generated a dual C-terminal FLAG-HA epitope tagged allele (Figure S1a), termed *Cntd1^FH^*. *Cntd1^FH/FH^* male mice are indistinguishable from wildtype littermates with respect to fertility, testis weight, sperm count, testis morphology, MLH1 accumulation in pachynema, and chiasmata formation (Figure S1).

Annotation of the *Cntd1* genomic locus describes a 7-exon gene encoding a 334 amino acid protein with a calculated molecular weight of approximately 40 kDa for the full-length, epitope-tagged form (Figure 1a, S2a) (NCBI ID: NM_026562). Western blotting of whole testis extracts from *Cntd1^+/+^*, *Cntd1^FH/+^*and *Cntd1^FH/FH^* adult matched littermate males demonstrated presence of the protein encoded by the allele specifically in epitope-tagged mice, but revealed a smaller than expected band at ∼30 kDa (Figure 1b, arrow). Prior characterization of the *Cntd1* locus described the use of a start codon near the beginning of exon 3 (NCBI ID: DS033671), which would produce a 27.5 kDa endogenous protein and 29.7 kDa FLAG-HA-tagged protein, matching the size we observe by western blot (Figure 1a, 1b, S1a). We term this protein product “CNTD1 short form”. Since a non-specific band around the correct predicted size (40 kDa) for CNTD1^FH^ is observed, we undertook immunoprecipitation (IP) followed by western blotting to determine if a form of *Cntd1^FH^* exists that is obscured by this non-specific band. We observe enrichment of the smaller 30 kDa band in the IP fraction (Figure S1b, arrow), loss of the non-specific band around 40 kDa, and no additional specific bands (Figure S2b).

**Figure 1:**
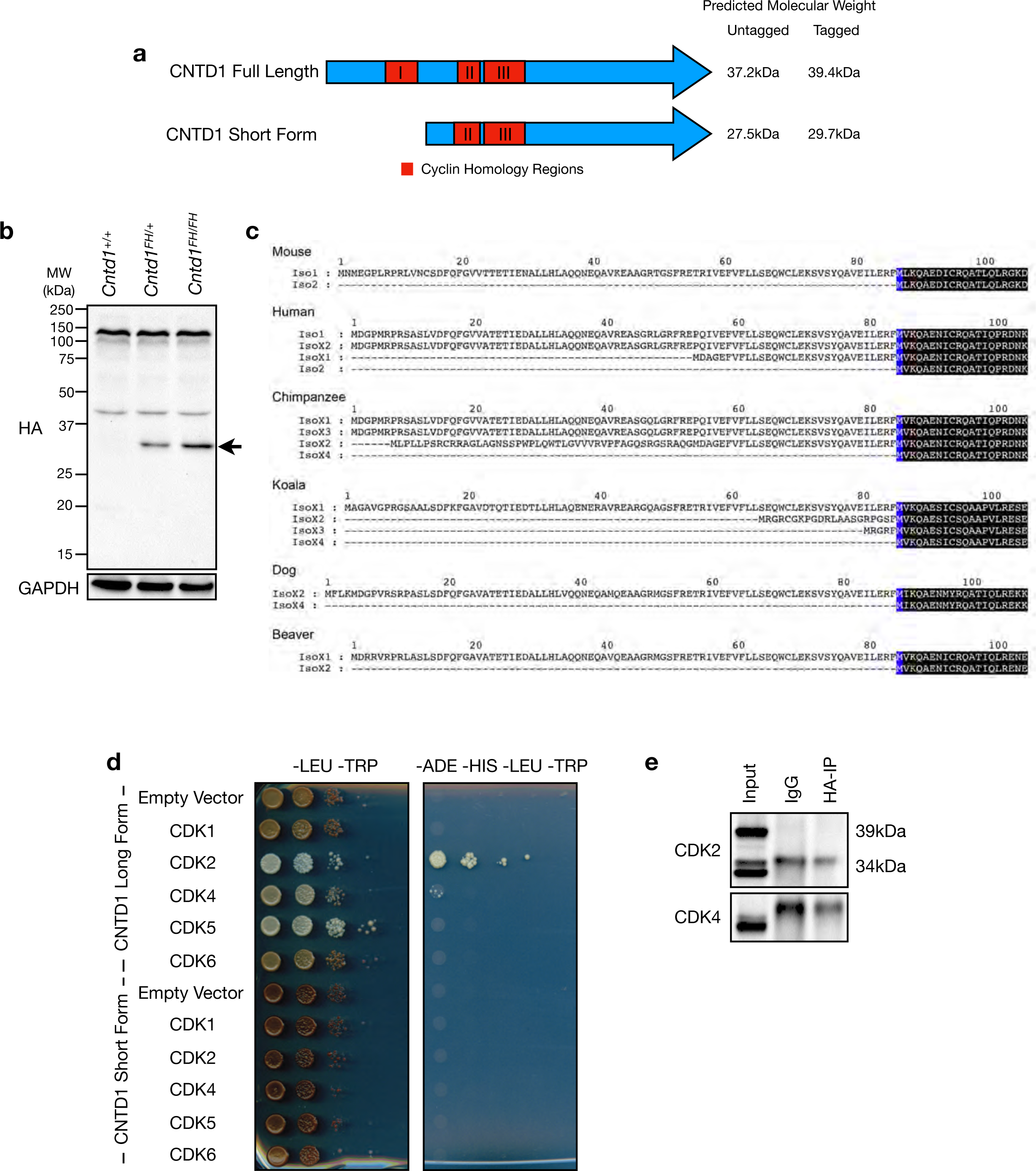
CNTD1^FH^ protein is smaller than expected, lacking an N-terminal cyclin homology domain required for CDK interaction. (a) To-scale schematic of annotated full-length CNTD1 and CNTD1 short form including predicted cyclin homology regions (red). (b) Western blot using antibodies against the HA epitope and GAPDH from testis lysate of *Cntd1^+/+^, Cntd1^FH/+^* and *Cntd1^FH/FH^* mice. Arrow indicates CNTD1 specific band. (c) N-terminal splice variants of mammalian CNTD1 homologs. Sequence alignments carried out using Clustal Omega (Sievers et al., 2011). Blue coloring denotes the conserved starting methionine of the short form in each homolog with the immediate downstream amino acid sequences that follow colored black. NCBI database IDs for annotated isoform sequences are as follows: Mus musculus (mouse), Iso1: NP_080838.1, Iso2: EDL01034.1; Homo sapiens (human), Iso1: NP_775749.2, IsoX2: XP_006721753.1, IsoX1: XP_005257100.2, Iso2: NP_001317151.1; Pan troglodytes (chimpanzee), IsoX1: XP_009430406, IsoX3, XP_016787256.1, IsoX2: XP_016787255.1, IsoX4: XP_024206054.1; Phascolarctos cinereus (koala), IsoX1: XP_020828270.1, IsoX2: XP_020828271.1, IsoX3: XP_020828272.1, IsoX4: XP_020828273.1; Canis familiaris (dog), IsoX2: XP_022278673.1, IsoX4: XP_022278675.1; Castor canadensis (American beaver), IsoX1: XP_020020731.1, IsoX2: XP_020020732.1. In mouse, Iso1 and Iso2 is equivalent to annotated full-length CNTD1 and CNTD1 short form respectively. Blue ‘M’ indicates conserved methionine. (d) Yeast two-hybrid using Clontech Y2H gold system, with annotated full-length CNTD1 and CNTD1 short form expressed as bait proteins with CDKs expressed as prey proteins, grown on control (-LEU-TRP) and selective plates (-ADE-HIS-LEU-TRP). (e) Western blot using antibodies against CDK2 and CDK4 from anti-HA immunoprecipitation of *Cntd1^FH/FH^*testis lysate.

We considered the possibility that standard sonication for protein extraction might not sufficiently liberate proteins associated with dense chromatin (Tran and Schimenti, Personal Communication). To determine if this might explain the absence of a native 40 kDa CNTD1 isoform, we undertook an extended sonication technique using protein from numerous tissues, but were unable to identify additional protein bands by western blot beyond those observed with routine sonication protocols (Figure S2c).

We also considered the possibility that the CNTD1 full-length protein may migrate at a speed that reflected a smaller molecular weight location than predicted. However, when we expressed constructs encoding the FLAG-HA-tagged full-length and short forms of CNTD1 in yeast, we detected both forms of the protein by western blotting using an anti-HA antibody (Figure S2e). Importantly, the mouse CNTD1 short form migrates at the 30 kDa size. Thus, we conclude that the sole CNTD1 protein variant found in mouse testis exists as a short form, 30 kDa protein.

### Comparative analysis of CNTD1 orthologs reveals N-terminal variability across species

The predominance of the mouse CNTD1 short form *in vivo* prompted us to examine the sequences of other orthologs. Sequence alignment shows strong conservation of CNTD1 across a diverse set of species with a few important caveats. First, we identify an internal methionine that is absolutely conserved throughout the entire family (Figure S3, blue). This methionine corresponds to the alternative start codon that produces the short form of CNTD1 in mouse (Figures 1b, S3). Second, we observe greater variability in the N-terminal portion of the protein upstream of this methionine (Figure 1b). This manifests not only as primary sequence differences between distant relatives like mouse and *C. elegans* COSA-1 (Figure S3), but also as splice variants within species that exhibit multiple annotated isoforms (Figure 1c). In these latter instances, splicing always modifies the N-terminus, leading to truncations that, at a minimum, constitute the short form of the protein. The koala ortholog exemplifies this pattern of variability (Figure 1c). It is important to note that one species, American alligator, has only a single short isoform that begins at the alternative methionine start site (Figure S3, bottom row). These *in silico* findings underscore the evolutionary conservation of the CNTD1 short form and suggest its broader biological importance.

### The CNTD1 N-terminus mediates interactions with cyclin-dependent kinases

CNTD1 has predicted homology to cyclin proteins across three regions: amino acids 57-88, 117-135 and 140-180 (Figure 1a). Truncation of the N-terminus in the CNTD1 short form removes cyclin homology region 1 (Figure 1a). Using yeast two-hybrid analysis, we investigated whether CNTD1 short form would associate with the CDKs that predominate in mouse spermatocytes: CDKs 1, 2, 4, 5, and 6^27–31^. The full-length mouse CNTD1 interacts with CDK2 and CDK4 in yeast two hybrid assays (Figure 1d, upper half). The CNTD1 short form, however, fails to interact with any of the tested CDKs (Figure 1d, lower half). To determine if CDK2 and CDK4 interactions with CNTD1 exist *in vivo,* we immunoprecipitated CNTD1^FH^ from mouse testis using anti-HA antibody and probed with antibodies against HA, as a control for immunoprecipitation, or CDK2 and CDK4. Immunoprecipitation with anti-HA antibodies failed to enrich for either CDK2 or CDK4 (Figure 1e), consistent with the conclusion that the CNTD1 short form is the predominant isoform in mouse and lacks the ability to interact with key prophase I CDKs.

### CNTD1^FH^ forms discrete foci in pachytene spermatocytes

We next undertook immunohistochemistry on testis sections from *Cntd1^+/+^* and *Cntd1^FH/FH^* matched-littermates. In testis sections from *Cntd1^FH/FH^* adults, we observe staining in primary spermatocytes using antibodies against the HA tag (Figure 2a, b, and S4a, b). The localization of CNTD1 appears largely nuclear and is restricted to cells in prophase I (Figure 2b, S4b). Some cytoplasmic localization is observed when staining intensity is increased by prolonged exposure to the DAB substrate (Figure S4b, square arrows), but staining remains heaviest in the nucleus (Figure S4b, arrows). No HA staining signal was observed in testis sections from wildtype littermate control males (Figure 2a and S4a).

**Figure 2:**
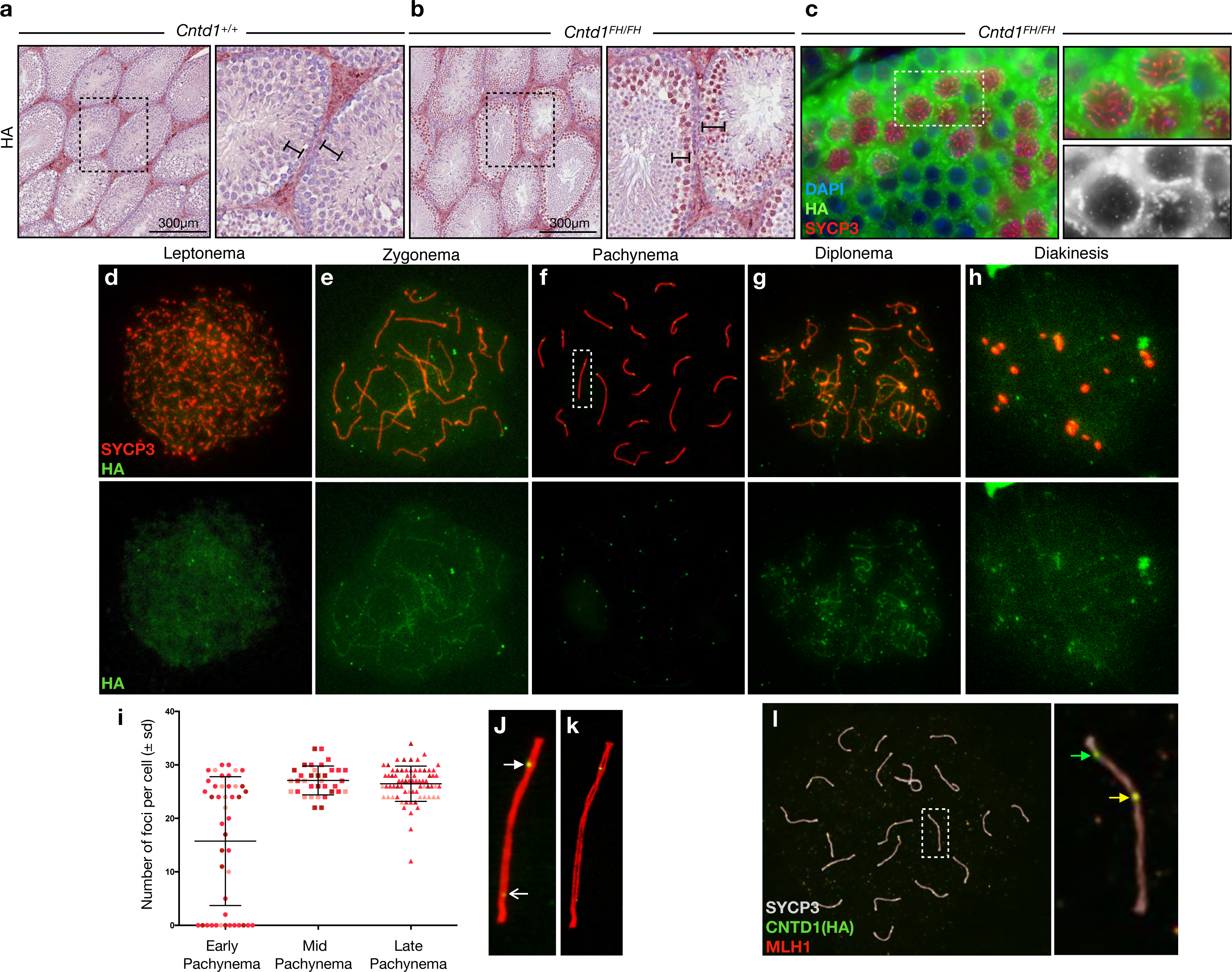
CNTD1^FH^ localizes to spermatocytes and forms discrete foci at pachytene. (a,b) Immunohistochemistry using anti-HA antibody on formalin fixed testis sections of *Cntd1^+/+^* and *Cntd1^FH/FH^* mice, imaged at 200x. Black bar indicates 300µm. (c) Immunofluorescence staining using anti-HA (green) and anti-SYCP3 (red) antibodies on formalin fixed testis sections of *Cntd1^+/+^* and *Cntd1^FH/FH^*mice, imaged at 200x. (d-h) Immunofluorescence staining on spread meiotic spermatocyte preparations from *Cntd1^FH/FH^*mice using antibodies against the HA epitope (green) and SYCP3 (red). Staging of spermatocytes defined by SYCP3 morphology. (i) Quantification of HA foci in pachynema staged spermatocytes of *Cntd1^FH/FH^* mice with mean and standard deviation plotted. Staging of early, middle, and late pachynema based upon the morphology of the XY chromosomes. (j) Enlarged image of dotted box in (f). Closed arrowhead indicates strong intensity focus; open arrowhead indicates low intensity focus. (k) Structural illumination microscopy rendering of (j). (l) Representative immunofluorescent staining of pachynema spermatocyte from *Cntd1^FH/FH^* testis using antibodies against MLH1 (red), HA (green) and SYCP3 (white). Right panel is enlarged insert of dotted box. Co-localization of HA and MLH1 identified by yellow arrow, HA independent foci identified by green arrow.

To further characterize the nuclear localization of CNTD1^FH^, we used immunofluorescence staining of fixed testis sections and on chromosome spread preparations. Both testis sections and spread meiotic spermatocytes showed discrete foci of CNTD1^FH^ at pachynema, distributed along the cores of the SC (Figure 2c-l). For chromosome spreads, we used co-localization with the SC lateral element protein SYCP3 to assess prophase I staging, providing a temporal profile of CNTD1^FH^ through prophase I. In leptonema, we observed diffuse nuclear staining of CNTD1^FH^, which accumulates along the SC in zygonema with some foci observed along the cores of the SC (Figure 2d,e).

Throughout pachynema, the diffuse nuclear staining of CNTD1^FH^ is no longer observed, being replaced instead by large discrete foci along the SC at a frequency reminiscent of MutLγ numbers (Figures 2f, j, k, l)^14, 15^. In early pachynema, localization of CNTD1^FH^ is observed in two distinct patterns: one population of cells has yet to accumulate any CNTD1^FH^ protein while a second population of cells shows increasing focus frequency (Figure 2i). By mid-pachynema, the focus frequency for CNTD1^FH^ protein is more homogenous, at 27 ± 2.7, and persisting at 26.5 ± 3.3 in late pachynema (Figure 2i). By diplonema, the diffuse nuclear staining pattern resumes but clear foci of HA signal remain associated with the SC, which disappear by diakinesis (Figure 2g-h).

From mid-pachynema onwards, the number of HA-marked CNTD1 foci is consistently higher than previously published focus frequencies for MLH1 and MLH3^14, 15^ and those reported herein (Figure S1f, 2i). Dual staining of CNTD1^FH^ and MLH1 reveals frequent presence of CNTD1^FH^ foci that do not co-localize with MLH1 foci but not *vice versa* (Figure 2l). Additionally, we observed two distinct intensity patterns for CNTD1^FH^ foci during pachynema: the majority of HA-stained foci are bright and robust, while approximately 1-5 foci per cell are of weaker intensity (Figure 2j). Importantly, increasing the exposure time for imaging of chromosome spreads does not reveal additional faint foci that are not included in our quantitation nor do we observe any non-specific HA-positive signal on chromosome spreads from wildtype males (Figure S4c-f).

### Mass Spectrometry of CNTD1^FH^-enriched STA-PUT fractions reveals interactions with RFC and SCF complexes

In order to define the temporal dynamics of CNTD1 appearance through prophase I relative to other key meiotic and cell cycle regulators, an improved gravitational cell separation (STA-PUT) strategy was devised to allow recovery of all stages of prophase I in a single procedure^32–34^. Meiotic chromosome spreads were prepared from 46 fractions and scored based upon SC morphology and presence of the phosphorylated histone variant, γH2AFX^1^.

The purity of STA-PUT enriched fractions for each germ cell stage reached 67% leptonema, 45% zygonema, 89% pachynema, 79% diplonema, and 98% sperm across different fractions (Figure 3). Western blotting performed against proteins from these cell fractions resulted in a dynamic profile of protein levels using a battery of antibodies against a range of meiosis-associated regulators (Figure 3), all compared to the dynamic expression of CNTD1 as detected by the HA tag. A thorough description of the protein profiles across the STA-PUT fractions is provided in the accompanying supplemental information (see Supplementary Material).

**Figure 3:**
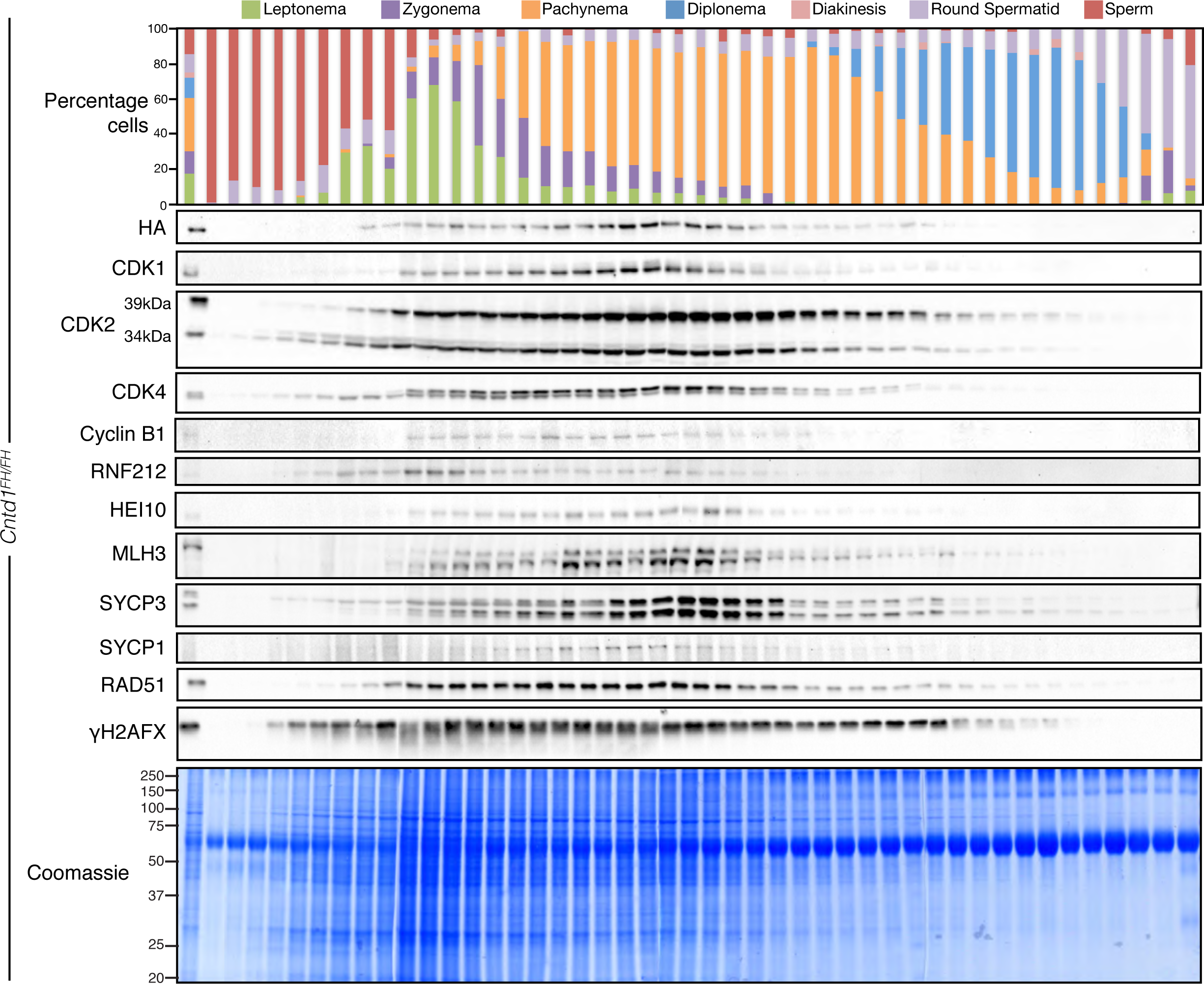
Gravitational cell separation (STA-PUT) allows isolation of prophase I stage specific fractions and reveals dynamic protein expression and post-translational modification. STA-PUT assay performed on *Cntd1^FH/FH^* testis single cell suspension. Top panel: Quantification of STA-PUT cell fraction composition based on a minimum of two hundred cells from spread preparations stained with antibodies against SYCP3 and γH2AFX. Western blots performed using the described antibodies against each STA-PUT fraction. Each western blot row is an amalgamation of four individual western blots performed and imaged under identical conditions.

Stage-matched prophase I cell fractions from *Cntd1^+/+^*and *Cntd1^FH/FH^* testes were pooled and used for immunoprecipitation against the HA epitope, followed by mass spectrometry (MS) to identify interacting proteins. A total of 588 proteins, as listed in Supplemental Table 1, were enriched in the CNTD1^FH^ sample compared with control sample, with 181 of these proteins identified by two or more peptides. Interestingly, given its localization pattern and function in crossover formation, we did not identify any proteins important for class I crossovers (MSH4, MSH5, MLH1, MLH3, RNF212, HEI10 or CDK2) nor did we identify any CDKs, consistent with the exclusive expression of *Cntd1* short form in mouse testis. Instead, the majority of CNTD1-interacting proteins fall into gene ontology groups involved in cellular and metabolic processes (Figure S6a). Given our interests in crossover designation, cell cycle progression, and the recent understanding of the role of post-translational modifications in driving these functions during meiosis ^35–37^, we chose to focus our attention on two specific complexes of interest. Both complexes involved at least two components that were shown in our MS data to interact with CNTD1. Firstly, CNTD1^FH^ specifically interacts with key regulators of meiotic processes and DNA repair, particularly components of the Replication Factor C (RFC) complex, which in somatic cells acts in the recruitment/activation of the DNA mismatch repair pathway^26, 38, 39^. CNTD1^FH^ also interacts with a large number of proteins belonging to the ubiquitylation pathway (Figure S6b), including components of the SKP1-Cullin-F-Box (SCF; reviewed in^40^), namely the E2 Conjugating enzyme CDC34 and the novel E3 ubiquitin ligase FBXW9 (Figure S6b).

### Replication Factor C (RFC) Complex

Replication factor C is a pentameric complex, consisting of RFC1 through RFC5, that loads PCNA during DNA replication^41^. RFC and PCNA have been implicated in activation of MutL endonucleases^26, 38, 39, 42^. Our mass spectrometry data revealed CNTD1 interaction with RFC3 and RFC4 (Supplemental table 1) and was confirmed by anti-HA-immunoprecipitation followed by western blot using anti-RFC3 and RFC4 antibodies (Figure 4a). Western blot analysis on whole testis lysate from *Cntd1^+/+^* and *Cntd1^-/-^* mice demonstrated no difference in the total protein levels of RFC3, RFC4, or PCNA (Figure 4b). In STA-PUT fractions from *Cntd1^+/+^* testes, we detected RFC3 and RFC4 protein in leptotene-enriched fractions through to pachytene-enriched fractions with PCNA signal observed through to diplotene containing cells (Figure 4d). In STA-PUT fractions from *Cntd1^-/-^* testes, RFC3 and RFC4 protein levels were decreased in lanes representing mostly pachytene cells, compared to similar lanes in WBs from STA-PUT-sorted *Cntd1^+/+^* fractions (Figure 5l).

**Figure 4:**
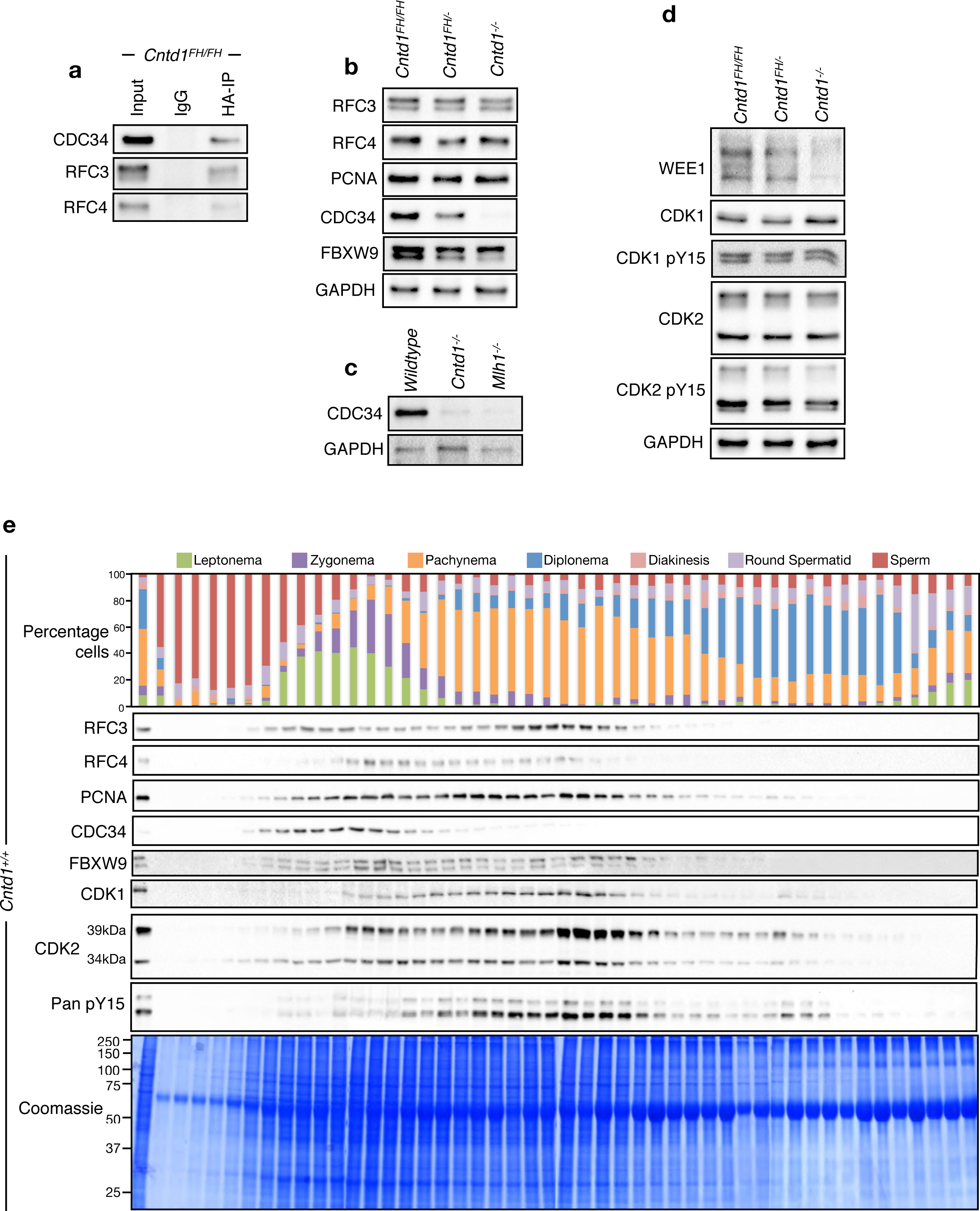
CNTD1 mutation leads to a decrease in CDC34 levels and lacks nuclear FBXW9 protein compared to wildtype. (a, b) Western blotting using antibodies against CNTD1^FH^ interacting proteins from immunoprecipitated material using anti-HA antibody from *Cntd1^FH/FH^*mice (a) and (b) from testis lysate of *Cntd1^FH/FH^, Cntd1^FH/-^*and *Cntd1^-/-^* mice. (c) Western blotting using antibodies against CDC34 and GAPDH on lysate from whole testis, *Wildtype, Cntd1^-/-^*and *Mlh1^-/-^*mice. (d) Western blotting against downstream SCF complex targets using antibodies against WEE1, CDK1, CDK1 pY15, CDK2, CDK2 pY15 and GAPDH. (e) STA-PUT assay performed on *Cntd1^FH/FH^* testis single cell suspension showing expression of CNTD1^FH^ interacting factors and downstream interactors relative to prophase I stage. Top panel: Quantification of STA-PUT cell fraction composition based on a minimum of two hundred cells from spread preparations stained with antibodies against SYCP3 and γH2AFX. Western blots performed using the described antibodies against each STA-PUT fraction. Each western blot row is an amalgamation of four individual western blots performed and imaged under identical conditions.

**Figure 5:**
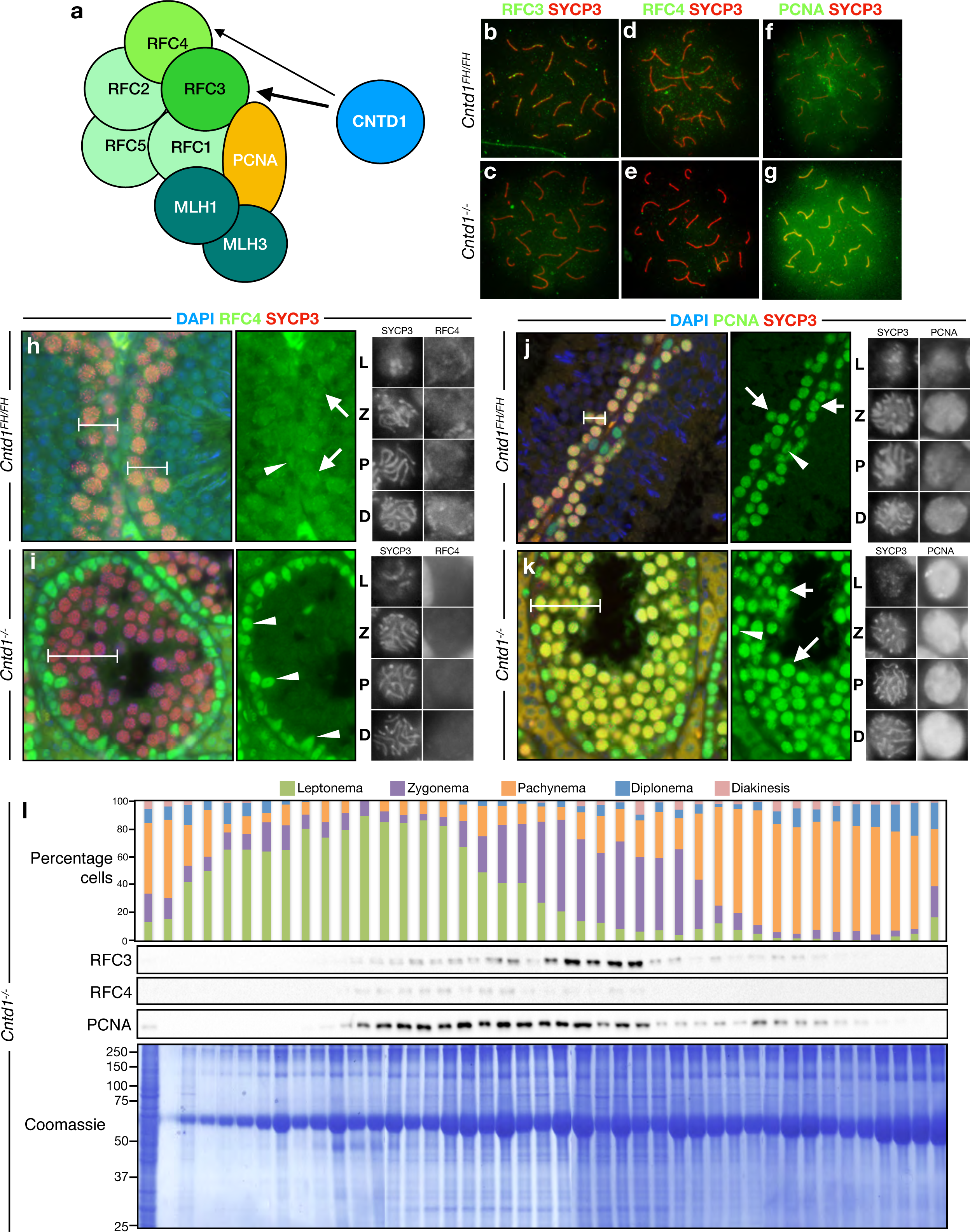
CNTD1 mutation causes a decrease in nuclear RFC4 signal and persistent increased nuclear signal of PCNA in prophase I cells. (a) Cartoon model depicting CNTD1 interactions (arrows) with components of the RFC complex and PCNA. Hypothesized interaction between RFC-PCNA and MLH1/MLH3 (MutLγ) based upon *in vitro* experiments requiring RFC and PCNA for activation of other described MutL complexes. (b - g) Immunofluorescence staining of spread meiotic spermatocytes from *Cntd1^FH/FH^*and *Cntd1^-/-^* mice using antibodies against RFC3, RFC4, PCNA and SYCP3. (h-i) Immunofluorescence staining using antibodies against RFC4 (green) and SYCP3 (red) on formalin fixed testis sections of *Cntd1^FH/FH^*(b) and *Cntd1^-/-^* (c) mice, imaged at 200x. White bars indicate prophase I cells. Arrowheads indicate spermatogonia and arrows indicate spermatocytes. Right hand panels showing representative images of prophase I stages from enlarged sections of (h) and (i) respectively, staged by the morphology of the synaptonemal complex from SYCP3 staining. (j, k) Immunofluorescence staining using antibodies against PCNA (green) and SYCP3 (red) on formalin fixed testis sections of *Cntd1^FH/FH^* (j) and *Cntd1^-/-^* (k) mice, imaged at 200x. White bars indicate prophase I cells. Arrowheads indicate spermatogonia and arrows indicate spermatocytes. Right hand panels showing representative images of prophase I stages from enlarged sections of (j) and (k) respectively, staged by the morphology of the synaptonemal complex from SYCP3 staining. (l) STA-PUT assay performed on *Cntd1^-/-^* testis single cell suspension showing expression of RFC3, RFC4 and PCNA relative to prophase I stage. Top panel: Quantification of STA-PUT cell fraction composition based on a minimum of two hundred cells from spread preparations stained with antibodies against SYCP3 and γH2AFX. Western blots performed using the described antibodies against each STA-PUT fraction. Each western blot row is an amalgamation of four individual western blots performed and imaged under identical conditions

Immunofluorescence staining on spread meiotic spermatocytes using antibodies against RFC3 and RFC4 revealed faint nuclear staining and focus formation along regions of the synaptonemal complex in pachytene-staged *Cntd1^FH/FH^* cells. In pachytene spermatocytes from *Cntd1^-/-^* males, the overall nuclear staining of RFC3 and RFC4 was reduced in addition to specific reductions in the number of RFC3/4foci forming along autosomal synaptonemal complexes compared with that observed in *Cntd1^FH/FH^* cells (Figure 5b-e). By contrast, PCNA immunofluorescence staining in pachytene spermatocytes from *Cntd1^FH/FH^* males revealed diffuse nuclear staining in addition to PCNA-loaded foci, some reminiscent of the patterning observed when staining for class I crossover proteins such as MLH1 and MLH3 (Figure 5f). By contrast, PCNA staining in pachytene spermatocytes from *Cntd1^-/-^* males was more intense across the entire nucleus, including the signal observed along the length of the synaptonemal complex (Figure 5g).

Immunolocalization of RFC4 protein in testis sections revealed robust RFC4 signal within the nuclei of spermatogonia and primary spermatocytes in wildtype testes (Figure 5a-b), with a steadily increasing signal for RFC4 as prophase I progresses (Figure 5b). In testis sections from *Cntd1^-/-^* males, we observe persistent, strong staining of RFC4 in spermatogonia (Figure 5c) but decreased staining in the primary spermatocytes at all stages of prophase I relative to that observed in *Cntd1^+/+^* (Figure 5c).

PCNA staining of *Cntd1^+/+^* testis sections revealed nuclear staining in occasional spermatogonia and all primary spermatocytes (Figure 5d). Similar staining is observed in primary spermatocytes throughout the *Cntd1^-/-^* tubule, with an apparent total increased intensity of PCNA signal due to the higher proportion of spermatocytes compared to that observed in adult *Cntd1^+/+^* testis (Figure 5e). However, the staining intensity in individual sub-stages of prophase I appears more robust and more persistent in spermatocytes from *Cntd1^-/-^* males than in wildtype controls (Figure 5d-e, right-hand grey panels). More importantly, these results demonstrate robust localization of RFC and PCNA in late stage primary spermatocytes, distinct from their expected staining in cells undergoing DNA replication (e.g. proliferating spermatogonia). In this regard, RFC and PCNA appear to play a role in DSB repair during meiosis that is analogous to their role in DNA repair in somatic cells ^41^.

### SKP1-Cullin-F-Box (SCF) Complex

The SKP1-Cullin-F-Box protein complex (SCF) is the most widely characterized of the superfamily of Cullin/RING ubiquitin ligases (CRL)^43^. The SCF core complex consists of RBX1, SKP1, Cullin 1, and a variable E3 ligase F-Box protein that collectively work together with E1 (activating) and E2 (conjugating) enzymes to define ubiquitylation substrates (Figure 6a)^44^. Our mass spectrometry data revealed that CNTD1 interacts with 22 components of the ubiquitylation/de-ubiquitylation family (Figure S6b and Supplemental table 1), including the E2 enzyme CDC34 and a novel E3 ligase F-Box protein FBXW9, both of which are SCF components.

**Figure 6:**
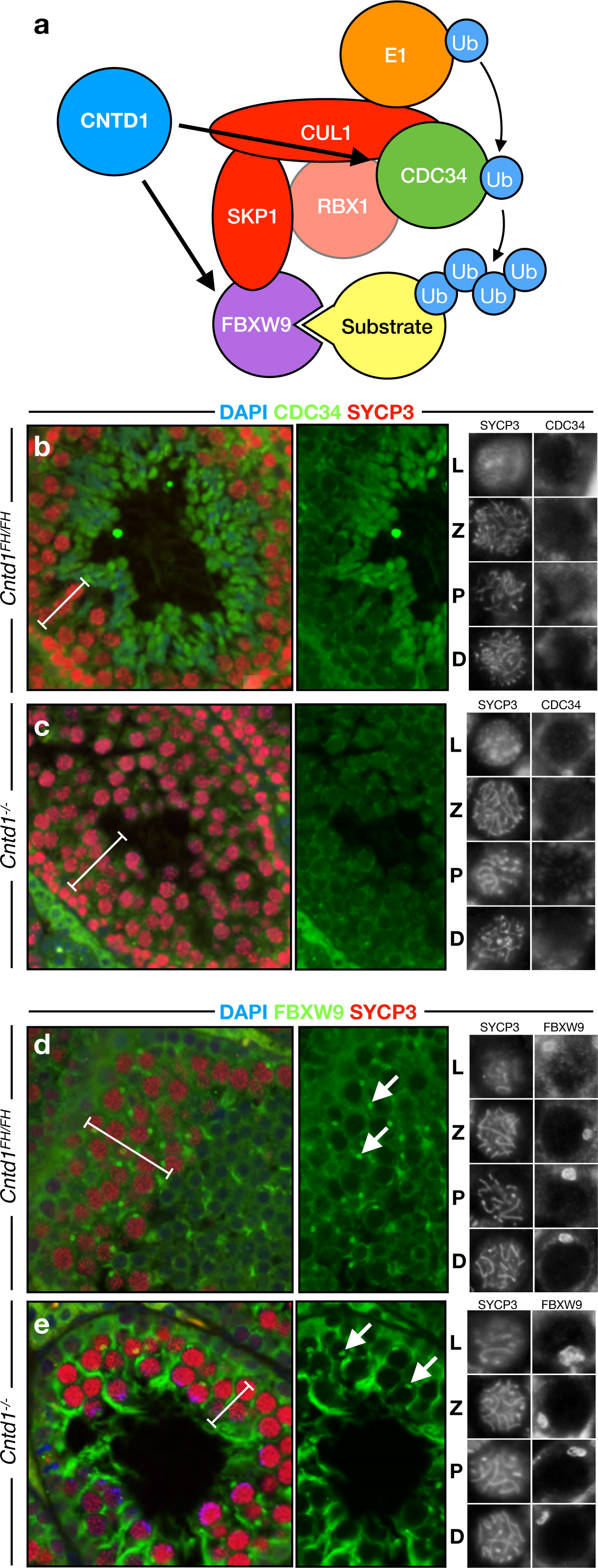
CDC34 levels are decreased in CNTD1 mutants and FBXW9 forms cytoplasmic protein aggregates that persist through prophase I. (a) Cartoon model depicting CNTD1 interactions (arrows) with components of the SCF complex. (b, c) Immunofluorescence staining using antibodies against CDC34 (green) and SYCP3 (red) on formalin fixed testis sections of *Cntd1^FH/FH^* (b) and *Cntd1^-/-^* (c) mice, imaged at 200x. White bars indicate prophase I cells. Right hand panels showing representative images of prophase I stages from enlarged sections of (b) and (c) respectively, staged by the morphology of the synaptonemal complex from SYCP3 staining. (d, e) Immunofluorescence staining using antibodies against FBXW9 (green) and SYCP3 (red) on formalin fixed testis sections of *Cntd1^FH/FH^*(d) and *Cntd1^-/-^* (e) mice, imaged at 200x. White bars indicate prophase I cells. Arrows indicate cytoplasmic protein aggregates. Right hand panels showing representative images of prophase I stages from enlarged sections of (d) and (e) respectively, staged by the morphology of the synaptonemal complex from SYCP3 staining.

The distribution of CDC34 and FBXW9 protein localization across STA-PUT fractions from *Cntd1^FH/FH^* males revealed detectable protein from leptonema through pachynema (Figure 4e). Immunoprecipitation followed by western analysis confirmed the interactions between CNTD1^FH^ and CDC34 (Figure 4a) while Western blot analysis of SCF components in whole testis lysates revealed that CDC34 levels in *Cntd1^-/-^* mutants were drastically decreased compared with that of *Cntd1^FH/FH^* animals (Figure 4b). Given this direct interaction, we next asked whether CDC34 levels were similarly altered in crossover mutants for genes acting downstream of CNTD1. CDC34 protein levels were reduced in *Mlh1^-/-^* mice to a similar extent compared with the reduction observed in *Cntd1^-/-^* mice (Figure 4c). Therefore, the lack of CDC34 protein is a consequence of a failure to form a competent crossover complex.

Immunofluorescence staining on testis sections revealed the most prominent staining for CDC34 in round and elongating spermatids of *Cntd1^FH/FH^* males. *Cntd1^-/-^* males, lacking these post-meiotic cell types, failed to show significant CDC34 staining (Figure 6b-c). In wildtype spermatocytes, CDC34 signal is restricted to the cytoplasm throughout prophase I (Figure 6b) and this localization is slightly reduced in spermatocytes from *Cntd1^-/-^*males (Figure 6c). By contrast, staining for FBXW9 revealed spermatocyte-specific aggregation of protein within the cytoplasm close to the nucleus in *Cntd1^FH/FH^* and *Cntd1^-/-^*spermatocytes (Figure 6d-e). In spermatocytes from *Cntd1^FH/FH^* and *Cntd1^-/-^* mice, FBXW9 protein appears restricted to the cytoplasm and of equivalent intensity (Figures 6d-e).

### Targets of SCF are drivers of cell cycle progression and crossover regulation during prophase I

CDC34, as part of SCF, has previously been shown to target the WEE1 kinase for degradation^45^. WEE1 functions as a mitotic inhibitor^45–49^, specifically phosphorylating the CDK1 component of maturation promoting factor (MPF) to prevent premature metaphase I entry^49–51^. WEE1 degradation thus allows MPF activation to facilitate cell cycle progression. Accordingly, WEE1 staining in fixed testis sections revealed strong cytoplasmic and diffuse faint nuclear localization in prophase I spermatocytes from *Cntd1^FH/FH^* males, with the nuclear localization disappearing by diplonema (Figure 7c).

**Figure 7:**
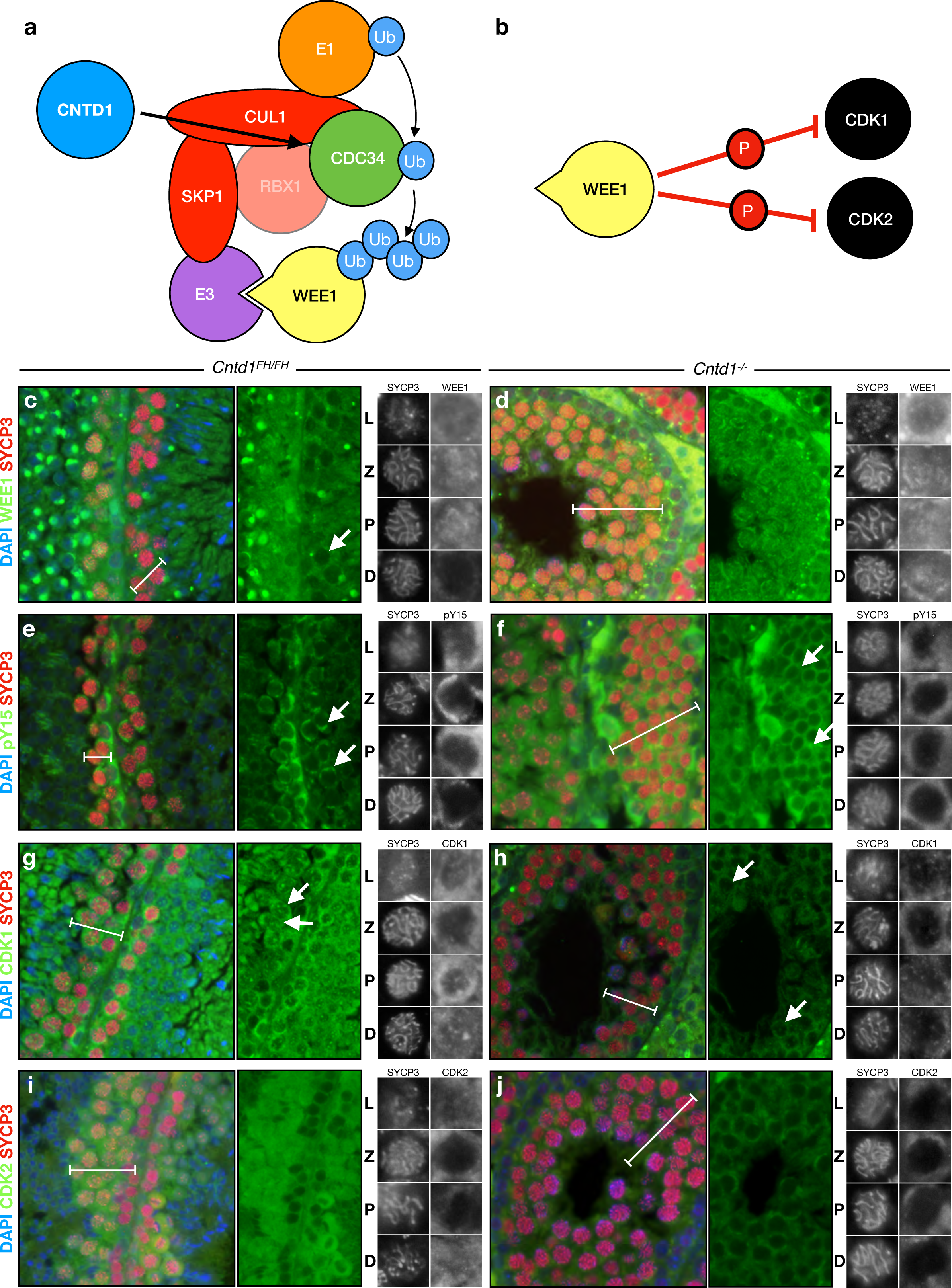
CNTD1 mutation leads to persistent nuclear WEE1 signal, increased cytoplasmic inhibitory phosphorylation on CDKs, and restricted CDK1 and CDK2 to the cytoplasm throughout prophase I. (a) Cartoon model depicting CNTD1 interactions (arrow) with WEE1 target of the CDC34 dependent SCF complex. (b) Cartoon depicting WEE1 inhibitory phosphorylation on CDK1 and CDK2. (c-j) Immunofluorescence staining using antibodies against representative antibodies on formalin fixed testis sections of *Cntd1^FH/FH^*(left column) and *Cntd1^-/-^* (right column), imaged at 200x. White bars indicate prophase I cells, arrow highlighting spermatocytes. Right hand panels showing representative images of prophase I stages from enlarged sections, staged by the morphology of the synaptonemal complex from SYCP3 staining.

Given the lack of post-spermatocyte cells within the seminiferous tubules of *Cntd1^-/-^* males and the decrease in testicular CDC34 protein in these animals, we hypothesized that regulation of WEE1 was defective in *Cntd1^-/-^* males. Indeed, total WEE1 was decreased in *Cntd1^-/-^* males compared with *Cntd1^FH/FH^*males whereas CDK1 and CDK2 protein levels were not significantly altered (Figure 4b). The nuclear localization of WEE1, however, is observed throughout prophase I and remains elevated at diplonema in spermatocytes of *Cntd1^-/-^*males rather than diminishing as in wildtype testes (Figure 7d).

WEE1 inhibits CDK1 and CDK2 activity by phosphorylation on tyrosine 15^52–55^, but phospho-specific antibodies against tyrosine 15 of CDK1 and CDK2 did not reveal any significant change in whole testis lysate from *Cntd1^-/-^* males compared with that of *Cntd1^FH/FH^* animals (Figure 4b). To asses stage specific changes in WEE1 phosphorylation, we stained testis sections from *Cntd1^FH/FH^* and *Cntd1^-/-^*males using a pan-CDK phospho-tyrosine 15 antibody (Figure 7e-f). We observed strong cytoplasmic staining of early stage prophase I cells in *Cntd1^FH/FH^* spermatocytes that is lost by diplonema (Figure 7e). By contrast, we observed retention of the cytoplasmic staining throughout spermatocytes at all stages of prophase I in the seminiferous epithelium of *Cntd1^-/-^* males (Figure 7f).

CDK1 and CDK2 staining in *Cntd1^FH/FH^*cells appear to be dynamic between the cytoplasm and nucleus throughout prophase I (Figure 7g, i). In early prophase I spermatocytes, CDK1 and CDK2 are distributed between both cellular compartments but by diplonema show strong nuclear staining. By contrast, staining of CDK1 and CDK2 in testis sections from *Cntd1^-/-^* males reveals much less intense nuclear signal with only faint cytoplasmic staining and no localization in the nucleus by diplonema (Figure 7h, j).

These data suggest that SCF components are misregulated in *Cntd1^-/-^* males relative to *Cntd1^FH/FH^*, leading to decreased CDC34 levels. This in turn results in increased WEE1 positive cells within the seminiferous epithelium of *Cntd1^-/-^* males coupled with a decrease in CDK1 and CDK2 staining in the nucleus of late prophase I cells and a persistence of phosphorylated (and thus inactivated) CDKs in the cytoplasm (Figure 7e-j). We postulate that the insufficient degradation of WEE1 arising from loss of CNTD1 and CDC34 directly contributes to the failure of cell cycle progression.

### *In vitro* inhibition of WEE1 kinase facilitates rapid progression into metaphase I and is a pre-requisite for spindle assembly checkpoint activation

Following prophase I, cells progress to metaphase I, the stage at which the spindle assembly checkpoint (SAC) monitors correct microtubule attachment before entry into anaphase I. Given our data, we postulated that spermatocytes from *Cntd1^-/-^* males fail to progress past prophase I to the metaphase-to-anaphase transition, and thus do not activate the SAC. To test this hypothesis, we cultured prophase I cells from *Cntd1^+/+^* and *Cntd1^-/-^* male mice for short periods of time, in the presence of Adavosertib (MK-1775) (0.25-10 µg/ml), a small molecule inhibitor that blocks WEE1 kinase activity, and/or Nocodazole (1-80 µg/ml), a microtubule polymerization inhibitor. As a readout for drug activity, we used western blot analysis to monitor levels of the anaphase I inhibitor MAD2L2 and inhibitory phospho-tyrosine 15 on CDK1 and CDK2 for SAC and WEE1 activity respectively.

Cells obtained from *Cntd1^-/-^* males showed a reduction in CDK1 and CDK2 inhibitory phosphorylation when treated with Adavosertib alone, while these same cells treated with Nocodazole alone failed to show any increase in the levels of MAD2L2 (Figure 8a,b). Only when both drugs were applied to these cells was any inhibition of anaphase I progression observed, reflected by an increase in MAD2L2 protein levels (Figure 8a,b). Accordingly, whole testis lysates from *Cntd1^-/-^* males showed a decrease in MAD2L2 protein levels compared to that of wildtype males (Figure 8c), indicative of a reduced number of cells completing prophase I.

**Figure 8:**
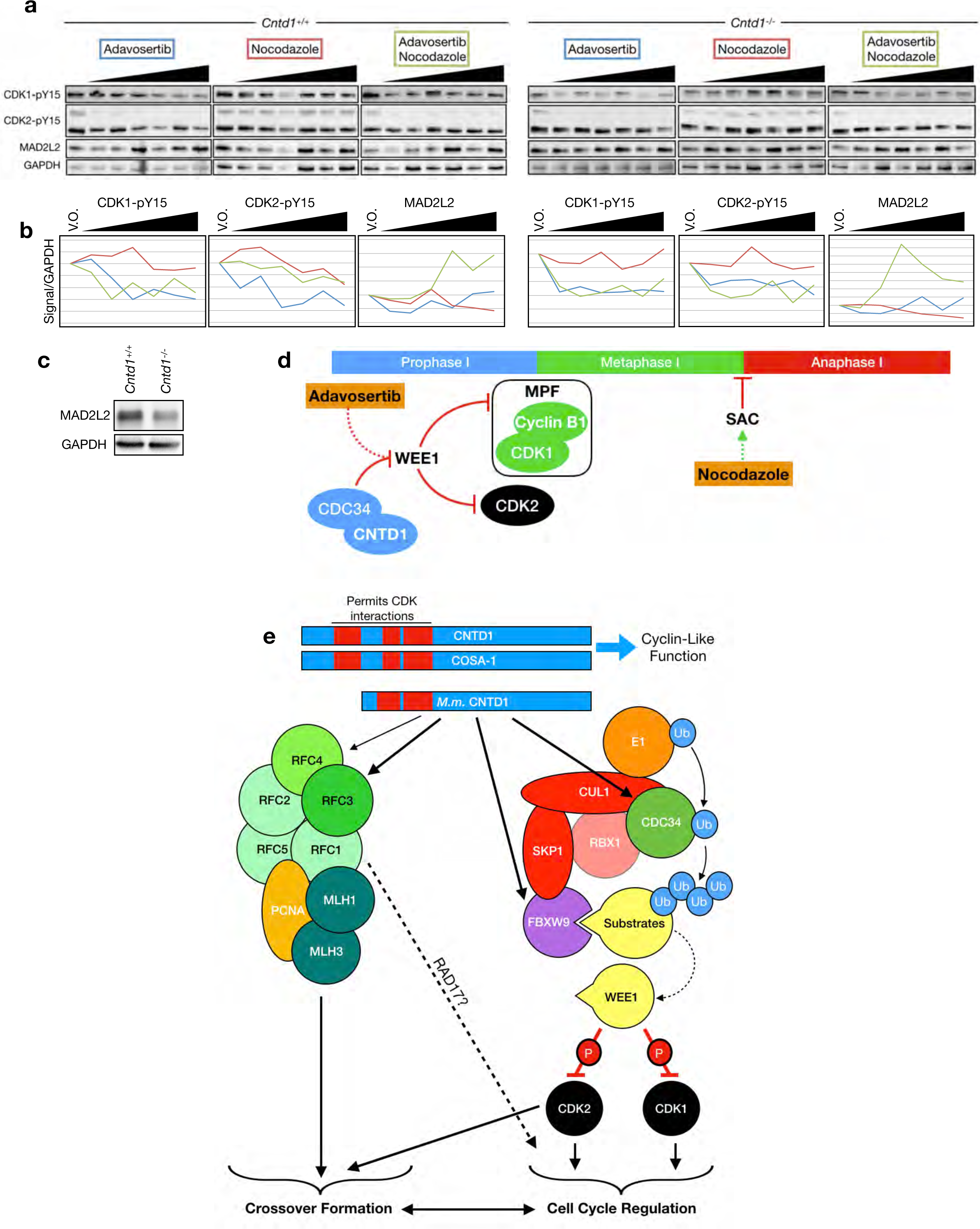
*In vitro* short-term cultures reveal a WEE1-dependent meiotic crossover checkpoint for progression into metaphase I. (a) Western blotting using antibodies against WEE1 inhibitory phosphorylation targets, CDK1-pY15 and CDK2-pY15, and anaphase inhibitor MAD2L2 following drug treatment of cultured spermatocytes from *Cntd1^+/+^ and Cntd1^-/-^* males. Drugs used were Adavosertib, Nocodazole and combined Adavosertib and Nocodazole treatment. Triangles indicate increasing drug concentrations, ranging from 0.25-10 µg/ml Adavosertib and 1-80 µg/ml Nocodazole. (b) Analysis of protein signal from western blotting in (a) divided by GAPDH signal, relative to the V.O. (vehicle only) control. (c) Western blotting using antibodies against MAD2L2 in whole testis lysates from *Cntd1^+/+^ and Cntd1^-/-^* males. (d) Cartoon depicting inhibitory phosphorylation of CDK1 and CDK2 by WEE1, which itself is degraded *in vivo* by CDC34 or inhibited by Adavosertib treatment. WEE1 degradation/inhibition allows activation of MPF and progression into metaphase I. The Spindle Assembly Checkpoint (SAC) monitors the correct attachment of microtubules and prevents the progression from metaphase I to anaphase I. Nocodazole inhibits microtubule formation and promotes/maintains the SAC. (d) Model of CNTD1 function. Mouse CNTD1 short form lacks the N-terminal cyclin homology domain that is required for direct interaction with CDKs in other organisms. CNTD1 interacts with components of two separate critical pathways required for the coordination of meiotic prophase I. CNTD1 interacts with components of the RFC complex, known to interact with PCNA, which in turn may activate MLH1/MLH3 for crossover formation. In addition, CNTD1 interacts with components of the SCF complex, which targets WEE1 for degradation allowing CDK1 and CDK2 activity and cell cycle progression. Cell cycle regulation may also occur through RAD17-RFC complex formation. Crossover formation is also driven by CDK2 at crossover sites.

To our surprise, cultured spermatocytes from wildtype males showed a similar requirement for both Adavosertib and Nocodazole treatment in order to elicit a MAD2L2 response (Figure 8a,b). These data indicate that inhibition of WEE1 is required for progression into metaphase, and that the treatment of wildtype spermatocytes with Adavosertib results in the massive and rapid induction of metaphase I entry, thus permitting progression through to anaphase I. Without Adavosertib, too few cells are naturally progressing into metaphase to be detected by MAD2L2 protein induction (Figure 8d).

## Discussion

Crossovers are essential for accurate segregation of homologous chromosomes during meiosis I, ensuring that gametes obtain the correct chromosome complement. Our previous work identified CNTD1, in addition to other crossover licensing and designation factors, as being critical for the formation of crossovers during prophase I in the mouse. CNTD1^FH^ foci are found in similar distribution and frequency to that of MutLγ, often co-localizing with this complex. We also observe multiple incidents of MLH1-independent CNTD1^FH^ foci at approximately 1-5 foci per cell, reminiscent of the number of crossovers expected to form through a MutLγ–independent mechanism ^19^. Thus, we propose that CNTD1 localizes to all nascent crossover sites, not just those that are destined to form class I, MutLγ-dependent crossovers. Importantly, we do not find any discrete CNTD1^FH^ signal prior to pachynema, suggesting CNTD1 accumulates at a time when class I and II crossovers are becoming designated.

Using CRISPR/Cas9, we generated an epitope-tagged allele of *Cntd1* that establishes a short form of CNTD1 – arising from translation of an internal start codon in exon 3 – as the predominant version present in the male mouse testis. This truncated protein lacks the first cyclin homology domain and is incapable of associating directly with any known CDKs. Our mass spectrometry and immunoprecipitation-western blot data, however, show that the CNTD1 short form can interact with regulators of CDK activity, thereby providing an indirect mechanism by which progression through the cell cycle can be modulated following appropriate crossing over. We cannot exclude the possibility that the CNTD1 short form maintains some cyclin-like functions since it still contains two cyclin homology domains with predicted structural similarity to cyclin A. Additionally, we cannot rule out that the full-length version of CNTD1 exists in low, undetectable levels or that it is more predominant in females. In testis from male mice, however, western blot analysis using an anti-HA antibody reveals a unique band migrating at a size that corresponds with the epitope-tagged CNTD1 short form, and this same antibody used on chromosome spread preparations localizes exclusively to sites that correspond with nascent crossovers.

Although the internal methionine start site used by mouse CNTD1 is absolutely conserved, the CNTD1 N-terminus shows a high degree of variability across species, with many organisms showing multiple isoforms in this region. These differences may reflect divergent CNTD1 functions, with some homologs retaining the ability to maintain direct CDK interactions and act as canonical cyclins while others must exert their effects indirectly. The presence of multiple isoforms may allow certain species to function via both mechanisms and respond to cellular and reproductive needs in a graded fashion.

Our mass spectrometry identifies interactions between CNTD1 and components of the RFC complex. While RFC and PCNA are required for the activity of human and yeast MutLα *in vitro* ^26, 38, 42^, yeast MutLγ displays activity independent of these proteins^56^. We observe decreased cellular signal of RFC4 but a stronger nuclear staining pattern of PCNA throughout prophase I in testis sections from *Cntd1* mutants. On meiotic spread preparations, PCNA localization along the SC increased in the absence CNTD1, coincident with decreased localization of RFC3/4. Previous characterization of *Cntd1^GT/GT^* male mice revealed normal prophase I progression prior to crossover formation as well as normal DSB formation and repair kinetics with no persistent DSBs observed ^25^. Early DSB repair kinetics also appear normal in *Cntd1^-/-^* animals. The persistent strong nuclear and SC staining of PCNA we observe within *Cntd1* null mutants is therefore not a consequence of failure to undergo appropriate early DNA repair events. This is further supported by diakinesis spreads that show decreased bivalent formation in *Cntd1^-/-^* animals but not fragmented chromosomes forming because of unrepaired DSBs. We propose that the increased PCNA staining in *Cntd1* mutants results from a failure to load CNTD1-RFC-PCNA complexes specifically at crossover sites and a lack of MutLγ, the presence of which in wildtype cells would stimulate crossover formation in the male mouse (Figure 8e).

Our results also indicate a robust interaction between CNTD1 and the SCF complex. Recent studies have highlighted the importance of ubiquitylation for key steps in germ cell development ^25, 37, 57, 58^, but the components of the ubiquitylation machinery acting in prophase I have not been fully elucidated. Here we describe the role of the SCF machinery in mammalian meiosis and the existence of a novel F-Box protein, FBXW9, presumably contained within this complex. CNTD1 interacts with several components of this complex, including the E2 conjugating enzyme CDC34. *Cntd1* null mutants, as well as mice lacking *Mlh1,* show a dramatic decrease in CDC34 protein levels. Though we have not identified the partner E3 ubiquitin ligase(s) that functions with CDC34, previous findings in *Xenopus* identified the cell cycle kinase WEE1 as a specific target of this adaptor^45^. WEE1 exerts inhibitory effects through phosphorylation of CDK1 and CDK2 and its removal enables mitotic cell cycle progression^49, 55, 59^. We find a similar mechanism functioning during meiotic prophase I in the male mouse. In this context, WEE1 phosphorylation of CDK1 maintains MPF in an inactive state and prevents cyclin B-CDK1-induced progression to metaphase. *Cntd1* null mutants show a retention of WEE1 in the nucleus throughout prophase I, continued inhibitory phosphorylation of its targets within the cytoplasm, and a consequent failure to localize CDK1 and CDK2 within the nucleus of late stage prophase I cells. These observations strongly argue that CNTD1 acts upstream of WEE1 through association with SCF and relieves the inhibitory effects of WEE1 phosphorylation, thereby coordinating the timing of cell cycle progression with proper crossover formation.

To clarify the connection between CNTD1, WEE1, and cell cycle progression, we explored how WEE1 inhibition might perturb the completion of prophase I. To test this in an *in vitro* short-term culture system, we utilized the small molecule WEE1 inhibitor, Adavosertib, in conjunction with the metaphase-to-anaphase inhibitor, nocodazole. Only with combined treatment using both drugs, were we able to observe inhibition of metaphase I progression, indicative of an induced SAC, and such a result was observed both in spermatocytes from wildtype and *Cntd1* mutant males. These results demonstrate, for the first time, the existence of a requirement for WEE1 degradation/inhibition to allow metaphase I entry and thus to facilitate progression to the metaphase-anaphase boundary (Figure 8d).

Collectively, our work defines two modes of action for CNTD1 in meiosis: simultaneously stimulating crossover formation through association with RFC while concomitantly regulating cell cycle progression through interactions with SCF and subsequent ubiquitylation of WEE1 (Figure 8e). Whilst these activities are distinct, the localization of both CNTD1^FH^ and CDK2 at crossover sites, together with the fact that CDK1 and CDK2 are both downstream targets of the SCF complex, supports the integrated regulation of the processes governing crossover formation and cell cycle progression by CNTD1. Our findings ultimately suggest that CNTD1 plays a critical role in ensuring that correct crossover levels are established before progressing into the first meiotic division. The function of CNTD1 in coordinating these processes suggests that CNTD1 acts as a stop/go regulator mechanism, monitoring crossover formation akin to a crossover-specific checkpoint, which may also be activated by cellular machinery downstream of CNTD1. CNTD1 interactions with RCF in the context of the RAD17-RFC complex, an established cell cycle checkpoint mediator^60^, could provide additional crosstalk and control of these pathways (Figure 8e). While our findings have immediate implications for the timing and control of prophase I in the male mouse, the dual nature of CNTD1 functions that we have uncovered here and the variable splicing apparent at the N-terminus implicate CNTD1 as a regulatory nexus throughout the animal kingdom.

## Acknowledgments

Research reported in this publication was supported by the Eunice Kennedy Shriver National Institute of Child Health & Human Development (NICHD) of the National Institutes of Health under Award Number R01HD041012 to P.E.C. Salary for S.G. was funded in part by a NICHD award to S.G. (K99HD092618). J.S.C. is a Meinig Family Investigator in the Life Sciences. Transgenic mice creation was supported in part by Empire State Stem Cell Fund Contract Number C024174. Imaging data were acquired through the Cornell University Biotechnology Resource Center, with NSF1428922 funding for the shared Zeiss Elyra super-resolution microscope. The content is solely the responsibility of the authors and does not necessarily represent the official views of the National Institutes of Health. We thank Dr. Attila Toth for sharing unpublished data, Lynn Dong for providing assistance with immunohistochemistry, Dr. Sheng Zhang and his team for performing Mass Spectrometry studies, and Rob Munroe and Chris Abratte of the Cornell Transgenic Mouse Core Facility for generation of new mouse lines. We also thank John Schimenti and Tina Tran for helpful conversations and Eric Alani, Michael Lichten, and Kathryn Grive for critical review of this manuscript.

## Author Contributions

S.G. and P.E.C conceived and designed the experiments, and analyzed the data; S.G. conducted the experiments; E.R.S. and J.S.C. performed computational alignment and analyzed data; S.G., J.S.C and P.E.C wrote the paper.

## Online Methods

### Mouse strains

All mouse alleles were maintained on a C57Bl6/J background, with at least six successive backcrosses. CRISPR/Cas9 genome edited mice were created initially on a FVB x C57B6 background before undergoing back crossing. All mice were maintained under strictly defined conditions of constant temperature and light:day cycles, with food and water *ad libitum*. Animal handling and procedures were performed following approval by the Cornell Institutional Animal Care and Use Committee, under protocol 2004-0063.

### Transgenic mice generation

*Cntd1-FLAG-HA:* CRISPR/Cas9 was used to insert the FLAG-HA epitope at the C terminus of the *Cntd1* locus. The RNA guide was generated by cloning the *Cntd1*specific sequence (GCCGCTTCCTCTAACACGTGA) in between the *BbsI* restriction sites of pX330, courtesy of Feng Zhang (Addgene #42230). Once integrated, the region including the gRNA scaffold was PCR amplified to include the T7 promoter. The PCR fragment was used to in an *in vitro* transcription reaction using the Ambion MEGAshortsript T7 Transcription Kit (AM1354).

Single-stranded DNA homology donor containing 78 nucleotides of homology upstream of the guide sequence, FLAG and HA epitopes separated by a leucine (generating a *HindIII* restriction site) and 66 nucleotides of homology downstream of the *Cntd1* stop codon was generated by IDT. gRNA, DNA donor and Cas9 mRNA was injected into 146 C57BL/6J x FVB F1 zygotes. 119 two-cell embryos were surgically implanted in 5 female recipients leading to the birth of 46 founder mice. PCR screening identified 7 founders with DNA integrated at the locus, 3 of which were the correct size. Sequencing revealed one founder mouse having the tags integrated in the correct frame. The founder mouse was backcrossed with C57BL/6J for four generations before being used for experiments described.

*Cntd1* null allele: CRISPR/Cas9 was used to generate a DNA double-strand break at the *Cntd1* locus and founder mice were screened for non-homologous end joining events generating a frame shift mutation. The gRNA was generated using long PCR primers incorporating the T7 promoter, targeting sequence and Cas9 scaffold followed by PCR and *in vitro* transcription (see above). Founders were screened for mutations by PCR of the *Cntd1* locus followed by sequencing. One founder carried an incorporation of 324 nucleotides of chromosome 12 at the targeted locus leading to disruption of the *Cntd1*open reading frame.

### Plasmid construction

*Yeast two-hybrid:* Testis cDNA was generated from Trizol extracted RNA using the Invitrogen SuperScript^®^ III First-Strand Synthesis System. CDKs 1, 2, 4, 5 and 6 were PCR amplified to include *NdeI* and *BamHI* restriction sites upstream and downstream respectively. The CDKs were cloned into Clontech yeast two-hybrid prey vector, pGADT7, using the Roche Rapid DNA Dephos & Ligation Kit. Annotated full-length CNTD1 and CNTD1 short form sequences were PCR amplified to include *NcoI* and *BamHI* restriction sites upstream and downstream respectively. The CNTD1 sequences were cloned into the Clontech yeast two-hybrid bait vector pGBKT7.

*CNTD1 expression constructs:* The yeast CEN/ARS region from pRS414 was PCR amplified, introducing restriction sites *SpeI* upstream and *EcoRV* downstream. This fragment was cloned into pFA6a-HphMX. The promoter of ADH1 from pGADT7 along with FLAG-HA epitope tagged CNTD1 full length and short forms from mouse testis cDNA was PCR amplified and cloned into the previously constructed vector using Gibson Assembly.

### Yeast two-hybrid assay

The yeast two-hybrid assay was performed as described by the manufacturer, Clontech. Briefly, combinations of bait and prey plasmids were transformed into the yeast two-hybrid reporter strain Y2HGold as described by Agatep *et al.* (1998) and plated onto –LEU –TRP selection plates. Positive colonies were re-streaked and tested for plasmids using PCR. Single colonies were inoculated into –LEU –TRP media and grown overnight. Colonies were normalized to an OD600 or 1.0 and four, 10 fold dilutions made. 10µl drops were placed on selection plates and quadruple dropout plates, which were incubated at 30°C for 3 days.

### Chromosome analysis and immunofluorescence

Chromosome spreads were made as previously described in Holloway et al. 2014. Briefly, decapsulated testis tubules were incubated in hypertonic elution buffer (30mM Tris pH7.2, 50mM sucrose, 17mM trisodium dehydrate, 5mM EDTA, 0.5mM DTT, 0.1mM PMSF, pH8.2-8.4) for one hour. Small sections of testis tubule were dissected in 100mM sucrose and spread onto 1% Paraformaldehyde, 0.15% Tiriton X coated slides and incubated in a humid chamber for 2.5hrs at room temperature. Slides were dried for 30minutes, washed in 1x PBS, 0.4% Photoflo (Kodak 1464510) and either stored at -80°C or stained. For staining, slides were washed in 1xPBS, 0.4% Photoflo for 10 minutes, followed by a 10 minute wash in 1x PBS, 0.1% Triton X and finally a 10 minute wash in 10% antibody dilution buffer (3% BSA, 10% Goat Serum, 0.0125% Triton X, 1 x PBS) 1x PBS. Antibodies used in this study are described in Supplemental table 1. Primary antibodies were diluted in 100% antibody dilution buffer, placed as a bubble on parafilm within a humid chamber, and the surface of the slide spread on the parafilm allowing the antibody to spread across the surface of the slide. Slides were incubated at 4°C overnight. Slides were washed in 1xPBS, 0.4% Photoflo for 10 minutes, followed by a 10 minute wash in 1x PBS, 0.1% Triton X and finally a 10 minute wash in 10% antibody dilution buffer. Secondary antibodies were diluted as the primary antibodies and spread in a similar fashion. Slides were incubated at 37°C for one hour. Slides were washed in 1xPBS, 0.4% Photoflo for 10 minutes, three times. Finally slides were left to dry, mounted using DAPI/antifade mix and either imaged or stored at 4°C for later imaging. Slides were imaged on a Zeiss Axiophot with Zen 2.0 software.

### Histology and immunofluorescence

Adult testes were dissected and incubated in either 10% neutral buffered formalin (for IHC) or Bouin’s fixative (for hematoxylin and eosin [H&E] staining) for 8hrs to overnight. Fixed testis was then washed 4 x in 70% ethanol, embed in paraffin and 0.5µM sections mounted on slides. For H&E staining, slides were rehydrated in safeclear followed by decreasing amounts of ethanol. Slides were then stained with hematoxylin followed by eosin then gradually dehydrated by incubation in increasing concentrations of ethanol. Finally, slides were mounted using permount and imaged on an Aperio Scanscope.

For immunofluorescence, slides were rehydrated as previously mentioned and then boiled in sodium citrate buffer (10mM Sodium Citrate, 0.05% Tween 20, pH 6.0) for 20 minutes. Following subsequent cooling, slides were blocked in blocking buffer (1 x PBST, 1% BSA, 3% Goat Serum) for an hour and primary antibody dilutions incubated on the sections overnight at 4°C. Slides were then washed, and incubated with fluorescence-conjugated secondary antibodies for one hour at 37°C, washed and then mounted using a DAPI/Antifade mix. Slides were imaged on a Zeiss Axiophot with Zen 2.0 software. Antibodies used in this study are described in Supplemental table 1.

### Sperm counts

Caudal epididymides of adult mice was transferred and dissected into sperm count media (4% BSA in 1 x PBS). Sample was incubated at 30°C for 30 minutes allowing sperm to swim out into the media. 1:10 dilutions were made in 10% neutral buffered formalin and stored at 4°C until counting on a haemocytometer.

### Spermatocyte diakinesis spread preparations

Spread diakinesis preparations were made as described in Holloway et al. 2014. Briefly, testis cells were liberated by manual dissection of tubules in 0.5% KCl. Multiple mixing and slow centrifugation followed by fixation in 30% methanol: 10% acetic acid: 0.05% chloroform. Cells were finally fixed in 30% methanol: 10% acetic acid and then pipette onto heated slides. Finally slides were stained in using Giemsa and imaged. Slides were imaged on a Zeiss Axiophot with Zen 2.0 software.

### Gravitational cell separation (STA-PUT)

Cells were separated using a protocol modified from Bryant et al. (2013). Testis extracts were dissected into 1 x Krebs buffer (Sigma K3753) supplemented with amino acids (Gibco 11130-051, Sigma M7145) and glutaMAX (Gibco 35050-061). Extract was incubated in a shaking water bath at 34°C, 150rpm in 2mg/ml Collagenase (Sigma C5138) in 1 x Krebs for 15 minutes. Following multiple rounds of centrifugation and washing in 1 x Krebs, the testis extract was re-suspended in 2.5mg/ml Trypsin (Sigma T0303), 200µg/ml DNAse (Sigma DN25) in 1 x Krebs and incubated in a shaking water bath at 34°C, 150rpm for 15 minutes. Extract was then centrifuged and washed multiple times in 1 x Krebs, followed by re-suspension in 0.5% BSA (Sigma A7906). Extract was then loaded into the STA-PUT apparatus along with different concentrations of BSA. Following loading of the BSA into the separating chamber along with the extract, the sample was allowed to sediment. Following sedimentation, cell fractions of were collected, centrifuged and washed in 1 x PBS. Small aliquots of each fraction were re-suspended in 50mM sucrose and incubated at room temperature for 20 minutes. 30µl drops of 1% Paraformaldehyde (EMS 19200) were placed on each well of an 8 well slide and the 50mM sucrose cell mix were added to each well. Slides were incubated in a humid chamber overnight, followed by drying, washing and staining against proteins defining stages of prophase I, in our case SYCP3 and γH2AX. The remaining extract was re-suspended in 1 x PBS Lysis buffer (1 x PBS, 0.01% NP-40, 5% Glycerol, 150mM NaCl, 1 x Roche cOmplete), sonicated and stored at -20°C for downstream applications.

### Protein extraction

Decapsulated testis extract was re-suspended into 1 x PBS Lysis buffer (1 x PBS, 0.01% NP-40, 5% Glycerol, 150mM NaCl, 1 x Roche cOmplete) and sonicated for 20 seconds at 22% amplitude in cycles of 0.4 seconds on and 0.2 seconds off.

### SDS-PAGE and Western Blotting

Protein samples were separated by SDS-PAGE on gels varying in percentage from 6-14% and transferred to methanol activated PVDF membranes using a Biorad Mini Trans-Blot Cell. Membranes were incubated in 5% BSA, 1 x TBST for 30minutes to 2 hours at room temperature whilst rotating at 60rpm. Membranes were incubated overnight in primary antibodies in 1 x TBST. Membranes were washed three times in 1 x TBST and subsequently incubated for one hour in secondary antibodies in 1 x TBST. Finally membranes were washed three times in 1 x TBST, developed using the ECL reagent and imaged using a Biorad ChemiDoc imager. Antibodies used in this study are described in Supplemental table 2.

### Colloidal Coomassie Staining

Protein extracts separated on SDS-PAGE gels were fixed in 40% methanol, 10% acetic acid for 30 minutes. Gels were stained using the Invitrogen Colloidal coomassie staining kit (LC6025) in 20% methanol, 20% stainer A, 5% stainer B and incubated at room temperature overnight whilst slowly rocking. Gels were subsequently washed in double distilled water and imaged.

### Image acquisition

Imaging was performed using a Zeiss Axiophot Z1 microscope attached to a cooled charge-coupled device (CCD) Black and White Camera (Zeiss McM). Images were captured and pseudo-colored using ZEN 2 software (Carl Zeiss AG, Oberkochen. Germany). Higher resolution images were acquired using an ELYRA 3D-Structured Illumination Super resolution Microscopy (3D-SIM) from Carl Zeiss with ZEN Black software (Carl Zeiss AG, Oberkochen. Germany). Images are shown as maximum intensity projections of z-stack images. To reconstruct high-resolution images, raw images were computationally processed by ZEN Black. Channel alignment was used to correct for chromatic shift. The brightness and contrast of images were adjusted using ImageJ (National Institutes of Health, USA).

### Mass spectrometry and protein identification

Mass spectrometry was performed in the Cornell University Proteomics and Mass Spectrometry facility. The nanoLC-MS/MS analysis was carried out using an Orbitrap Fusion (Thermo-Fisher Scientific, San Jose, CA) mass spectrometer equipped with a nanospray Flex Ion Source using high energy collision dissociation (HCD) and coupled with the UltiMate3000 RSLCnano (Dionex, Sunnyvale, CA). Each reconstituted sample was injected onto a PepMap C-18 RP nano trap column (3 µm, 100 µm × 20 mm, Dionex) with nanoViper Fittings at 20 μL/min flow rate for on-line desalting and then separated on a PepMap C-18 RP nano column (3 µm, 75 µm x 25 cm), and eluted in a 120 min gradient of 5% to 35% acetonitrile (ACN) in 0.1% formic acid at 300 nL/min. The instrument was operated in data-dependent acquisition (DDA) mode using FT mass analyzer for one survey MS scan for selecting precursor ions followed by 3 second “Top Speed” data-dependent HCD-MS/MS scans in Orbitrap analyzer for precursor peptides with 2-7 charged ions above a threshold ion count of 10,000 with normalized collision energy of 38.5%. For label-free protein analysis, one MS survey scan was followed by 3 second “Top Speed” data-dependent CID ion trap MS/MS scans with normalized collision energy of 30%. Dynamic exclusion parameters were set at 1 within 45s exclusion duration with ±10 *ppm* exclusion mass width. All data are acquired under Xcalibur 3.0 operation software and Orbitrap Fusion Tune 2.0 (Thermo-Fisher Scientific).

All MS and MS/MS raw spectra from each experiment were processed and searched using the Sequest HT search engine within the Proteome Discoverer 2.2 (PD2.2, Thermo). The default search settings used for relative protein quantitation and protein identification in PD2.2 searching software were: two mis-cleavage for full trypsin with fixed carbamidomethyl modification of cysteine and oxidation of methionine and demaidation of asparagine and glutamine and acetylation on N-terminal of protein were used as variable modifications. Identified peptides were filtered for maximum 1% false discovery rate (FDR) using the Percolator algorithm in PD 2.2. The relative label free quantification method within Proteome Discoverer 2.2 software was used to calculate the protein abundances. The intensity values of peptides, which were summed from the intensities values of the number of peptide spectrum matches (PSMs), were summed to represent the abundance of the proteins. For relative ratio between the two groups, here PGC female/male and Soma female/male, no normalization on total peptide amount for each sample was applied. Protein ratios are calculated based on pairwise ratio, where the median of all possible pairwise ratios calculated between replicates of all connected peptides.

### Short-term testis drug culture

Cultures were performed as described elsewhere ^61, 62^, with some modifications. Testis extracts were dissected into 1 x PBS and incubated in a shaking water bath at 34°C, 150rpm in 2 mg/ml Collagenase (Sigma C5138) in 1 x PBS for 15 minutes. Following multiple rounds of centrifugation and washing in 1 x PBS, the testis extract was re-suspended in 2.5 mg/ml Trypsin (Sigma T0303), 200 µg/ml DNAse (Sigma DN25) in 1 x PBS and incubated in a shaking water bath at 34°C, 150rpm for 15 minutes. Extract was then centrifuged and washed five times in 4 mL spermatocyte culture medium (SCM) (GIBCO DMEM without red phenol (21063-029), fetal calf serum (GIBCO 10082139), penicillin-streptomycin 1003 (GIBCO 15140-122); lactic acid (Sigma L13750, NaHCO3 9s8761) and sodium pyruvate 1003 (Sigma 11360-070). Cells were re-suspended in 600µl of SCM and placed in 12-well treated culture dishes (Corning #3513). Adavosertib (MedChemExpress HY-10993) was diluted in 10% DMSO and added at 0.25, 0.5, 1, 2, 5 and 10 µg/ml. Nocodazole (Sigma SML1665-1ML) was diluted in 10% DMSO and added at 1, 5, 10, 20, 50 and 80 µg/ml. Cultures were incubated at 37°C for 6 hours followed by protein extraction.

### Quantification and Statistical Analysis

Statistical analyses were performed using GraphPad Prism version 6.00 for Macintosh (GraphPad Software, San Diego California USA, www.graphpad.com). Specific analyses are described within the text and the corresponding figures. Mean values are all presented ± standard deviation (s.d.) and alpha value was established at 0.05. All statistical analyses performed using two-sided tests.

