## Supplemental Figures for "Cyclin N-Terminal Domain-Containing 1 (CNTD1) coordinates meiotic crossover formation with cell cycle progression in a cyclin-independent manner"

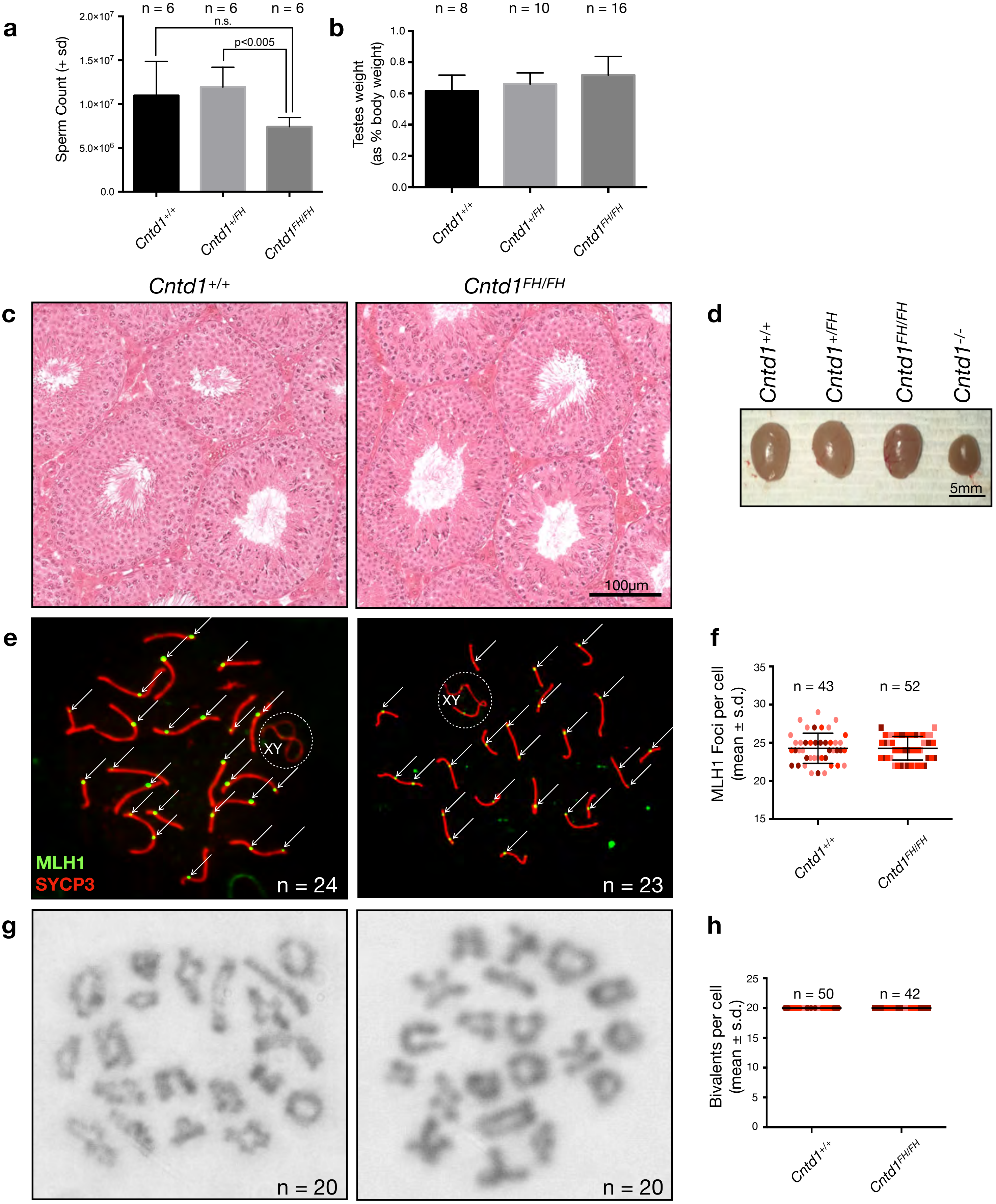
a
n = 6
n = 6
n = 6
n.s.
p<0.005
Sperm Count (+ sd)
2.0×10^7
1.5×10^7
1.0×10^7
5.0×10^6
0.0
Cntd1+/+
Cntd1+/FH
Cntd1FH/FH
b
n = 8
n = 10
n = 16
Testes weight
(as % body weight)
1.0
0.8
0.6
0.4
0.2
0.0
Cntd1+/+
Cntd1+/FH
Cntd1FH/FH
Cntd1+/+
Cntd1FH/FH
c
100μm
d
Cntd1+/+
Cntd1+/FH
Cntd1FH/FH
Cntd1-/-
5mm
e
XY
MLH1
SYCP3
n = 24
XY
n = 23
f
MLH1 Foci per cell
(mean ± s.d.)
35
30
25
20
15
n = 43
n = 52
Cntd1+/+
Cntd1FH/FH
g
n = 20
n = 20
h
Bivalents per cell
(mean ± s.d.)
25
20
15
10
5
0
n = 50
n = 42
Cntd1+/+
Cntd1FH/FH

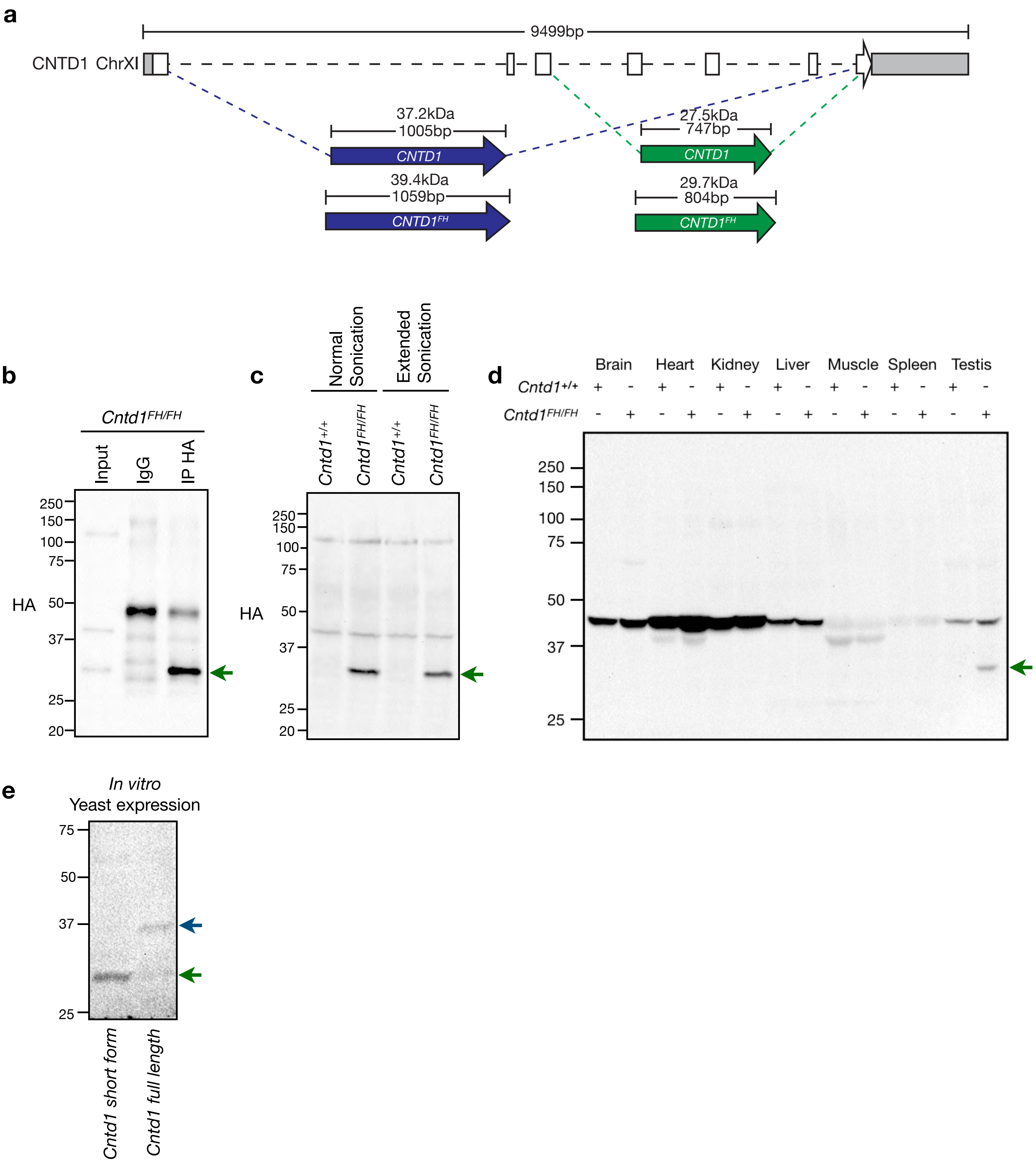
a
9499bp
CNTD1 ChrXI
37.2kDa
1005bp
CNTD1
39.4kDa
1059bp
CNTD1FH
27.5kDa
747bp
CNTD1
29.7kDa
804bp
CNTD1FH
b
Cntd1FH/FH
Input
IgG
IP HA
HA
250
150
100
75
50
37
25
20
c
Normal Sonication
Extended Sonication
Cntd1+/+
Cntd1FH/FH
Cntd1+/+
Cntd1FH/FH
HA
250
150
100
75
50
37
25
20
d
Brain
Heart
Kidney
Liver
Muscle
Spleen
Testis
Cntd1+/+
Cntd1FH/FH
250
150
100
75
50
37
25
e
In vitro
Yeast expression
75
50
37
25
Cntd1 short form
Cntd1 full length

Gray *et al*.
Figure S3

```
Mm_CNTD1      1 : ----------------------------------MNMEGPLRPR-LVNCSDFQFG---VVTTETIENA :  30
Hs_CNTD1      1 : ------------------------------------MDGPMRPR-SASLVDFQFG---VVATETIEDA :  28
Pt_CNTD1      1 : ------------------------------------MDGPMRPR-SASLVDFQFG---VVATETIEDA :  28
Pc_CNTD1      1 : ------------------------------------MAGAVGPRGSAALSDFKFG---AVDTQTIEDT :  29
Clf_CNTD1     1 : --------------------------------MFLKMDGPVRSR-PASLSDFQFG---AVATETIEDA :  32
Cc_CNTD1      1 : ------------------------------------MDRRVRPR-LASLSDFQFG---AVATETIEDA :  28
Ce_COSA-1     1 : MSSSRSHRKNTSTLGTPAVSAANQTVKNPNLKKNEPKSDNEPPKTLVSMEPDFYDPRGACHMIYWTDC :  68
Am_CNTD1      - : -------------------------------------------------------------------- :   -
                                                      m     p       df  g    v t tie

                                                                         *
Mm_CNTD1     31 : LLHLAQQNEQAVKEAAGRTGSFRETRIVEFVFLLSEQWCLEKSVSYQAVEILERFMLKQAEDICRQAT :  98
Hs_CNTD1     29 : LLHLAQQNEQAVREASGRLGRFREPQIVEFVFLLSEQWCLEKSVSYQAVEILERFMVKQAENICRQAT :  96
Pt_CNTD1     29 : LLHLAQQNEQAVREASGQLGRFREPQIVEFVFLLSEQWCLEKSVSYQAVEILERFMVKQAENICRQAT :  96
Pc_CNTD1     30 : LLHLAQENERAVREARGQAGSFRETRIVEFVFLLSEQWCLEKSVSYQAVEILERFMVKQAESICSQAA :  97
Clf_CNTD1    33 : LLHLVQQNEQAMQEAAGRMGSFRETRIVEFVFLLSEQWCLEKSVSYQAVEILERFMIKQAENMYRQAT : 100
Cc_CNTD1     29 : LLHLAQQNEQAVQEAAGRMGSFRETRIVEFVFLLSEQWCLEKSVSYQAVEILERFMVKQAENICRQAT :  96
Ce_COSA-1    69 : IAQMAVDIRERQNAANQSDFDFMKPKLVEYVFTVCVRLRLPNEVRFTAALILNSFMLRHLCSLHDFME : 136
Am_CNTD1      1 : -------------------------------------------------------MIKQVQEMYNSTE :  13
                   lh aq ne a  ea g  g fre   ve vfl seqwcleksvs qaveilerfM64qa  6  qa

Mm_CNTD1     99 : LQLRGK-DTELQSWRAMKEQLVNKFILRLVSCVQLASKLSFHYKIVSNITVLNFLQALGYVHTKEELL : 165
Hs_CNTD1     97 : IQPRDN-KRESQNWRALKQQLVNKFTLRLVSCVQLASKLSFRNKIISNITVLNFLQALGYLHTKEELL : 163
Pt_CNTD1     97 : IQPRDN-KRESQNWRALKQQLVNKFTLRLVSCVQLASKLSFRNKIISNITVLNFLQALGYLHTKEELL : 163
Pc_CNTD1     98 : PVLRESEKAELWSWRARKEELCSTFVLRLVSCVQLASKLSFHYKIVSNVTVLNFLQALGYLYTKEELL : 165
Clf_CNTD1   101 : IQLRE--KKEPQNWKALKEQLFNKFILRLVSCVQLASKLSFHYKIISNITVLNFLQSLGYVHTKEELL : 166
Cc_CNTD1     97 : IQLRENEKTDSQNWRALKEQLFNKFILHLVSCVQLASKLSFHYKIISNITVLNFLQALGYLYTKEELL : 164
Ce_COSA-1   137 : -RQEMSIQRKKREWENLESNMERQIPLRILTAIQISSKFHSYHDSLSSRQVVNTLRKIGLPYTISAVL : 203
Am_CNTD1     14 : ESGESGEQGRSHGWNFLKAQMHNMFMLRLVSCIQLASKLSFHYSIVNNNTVLKFLQSLDYSFTKQELL :  81
                     r         W a k  6 n f Lr663c6Q6aSKlsf  ki6sn tV6nfLq 6gy  Tk e6L

Mm_CNTD1    166 : ESELDILKSLNFQINLPTPLAYVEMLLEVLGYNG---CLVP---ATQLHATCLTLLDLVYLLHEPIYE : 227
Hs_CNTD1    164 : ESELDVLKSLNFRINLPTPLAYVETLLEVLGYNG---CLVP---AMRLHATCLTLLDLVYLLHEPIYE : 225
Pt_CNTD1    164 : ESELDVLKSLNFQINLPTPLAYVETLLEVLGYNG---CLVP---AMRLHATCLTLLDLVYLLHEPIYE : 225
Pc_CNTD1    166 : ESELAILKSLNFQINLPTPLAYVEMLLEVLGYNG---CLVS---VKRLHGTCLTLLDMVYLLHGPIYE : 227
Clf_CNTD1   167 : ESELDVLKSLNFQINLSTPLAYVEMLLEVLGYNG---CLVP---ATRLHATCLTLLDLVYLLREPIYE : 228
Cc_CNTD1    165 : ESELDVLKSLNFQINLPTPLAYVEMLLEVLGYNG---CLVP---ATQLHATCLTLLDLVYLLHEPIYE : 226
Ce_COSA-1   204 : ESEQRVFKLIGFKMPD-SPLDACEMALKVLTFTMKKRGMIDEEKYNDLWQHTLIVLDVCFINHIELYE : 270
Am_CNTD1     82 : ESELAILKGLHFQINLPTPLAYVELLLEVLGHNG---CLLS---LKQLHEMCVHLLDLTYLMRDVIYN : 143
                  ESEl 6lKs6nFq6nlp3PLayvE lLeVLg ng   c66       Lh tc6t6LD6v56 h p6Ye

Mm_CNTD1    228 : SLLRASIENSTPSQLQGEKFLSVKEDFMLLAVGIIAASAFIQNHECWSQVIGHLQSITGIASESIAEF : 295
Hs_CNTD1    226 : SLLRASIENSTPSQLQGEKFTSVKEDFMLLAVGIIAASAFIQNHECWSQVVGHLQSITGIALASIAEF : 293
Pt_CNTD1    226 : SLLRASIENSTPSQLQGEKFTSVKEDFMLLAVGIIAASAFIQNHECWSQVVGHLQSITGIALASIAEF : 293
Pc_CNTD1    228 : SLLRASIENCTPME-QGRKFVSVKEDFMLLAVGIIAASAFIQNQEWWGQVVEHLHSITSIATESIAEF : 294
Clf_CNTD1   229 : SLLRASIENPIPSQLQGEKFVSVKEDFMLLAVGIIAASAFIQNHECWSQVVGHLQSITGIALESIAEF : 296
Cc_CNTD1    227 : SLLKASIENSTPSQLQGEKFISVKEDFMLLAVGIIAASAFIQNHECWRQVVGHLQSITGIALESIAEF : 294
Ce_COSA-1   271 : RFIRKCPAICRTEERLN--ISKFKWDIQLLAAATVQTAYILLLG---TSQIANVSVIINNLLRCDNAY : 333
Am_CNTD1    144 : TLLKTSIENSTPNELQIAKFLSVKEDFMLLAAGIIGTSAFILNPEHWNQVVEHLNCITGITSQSVFEF : 211
                   l64asien tp 2lqg kf svKeDfmLLAvgi6aasaf6qn e w qv6 h6 sItgia  s ae5

Mm_CNTD1    296 : SYAILTHSVGANTPGPQQPVP-HKAARALRTAAAAASSNT : 334
Hs_CNTD1    294 : SYAILTHGVGANTPGRQQSIPPHLAARALK---TVASSNT : 330
Pt_CNTD1    294 : SYAILTHGVGANTPERQQSIPPHLAARALK---TVASSNT : 330
Pc_CNTD1    295 : SYAILTHGVGASTPVRQQPGLHHLAARPLR---AAATCNI : 331
Clf_CNTD1   297 : SYAILTHSVGANTPRRQQPVLPHLEARALR---AAASSNT : 333
Cc_CNTD1    295 : SYAILTHSVGANTPGQQQPVPPHLAARALR---AAASSNT : 331
Ce_COSA-1   334 : VEPLKQSIIELACAKKNESIPECSTSS------------- : 360
Am_CNTD1    212 : LYAILKHSLGTTPTGTN--------ARME----------- : 232
                  sya6lth 6ga tp  q     h  ar l      a  n
```

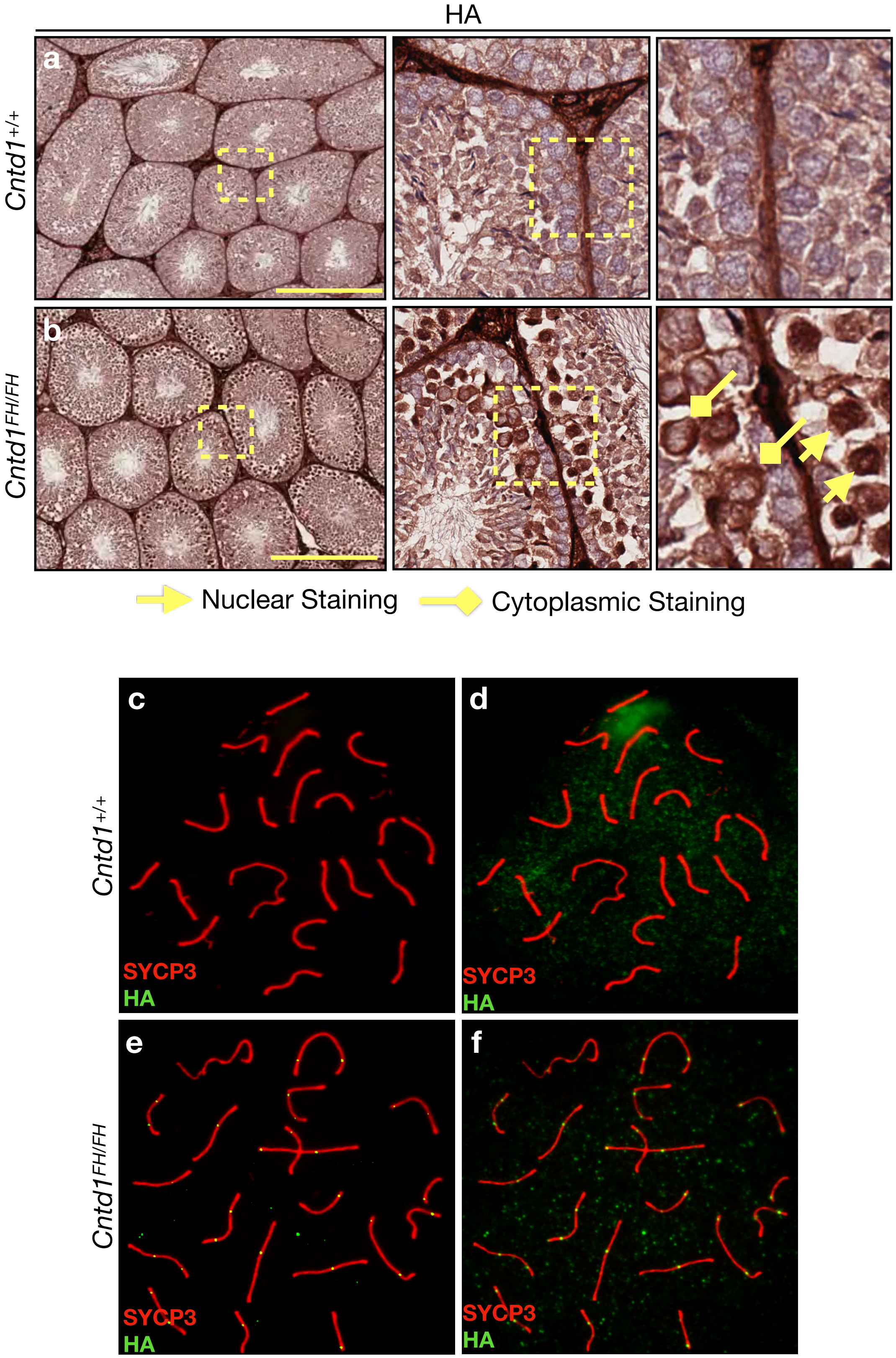
HA
a
Cntd1+/+
b
Cntd1FH/FH
Nuclear Staining
Cytoplasmic Staining
c
d
e
f
SYCP3
HA

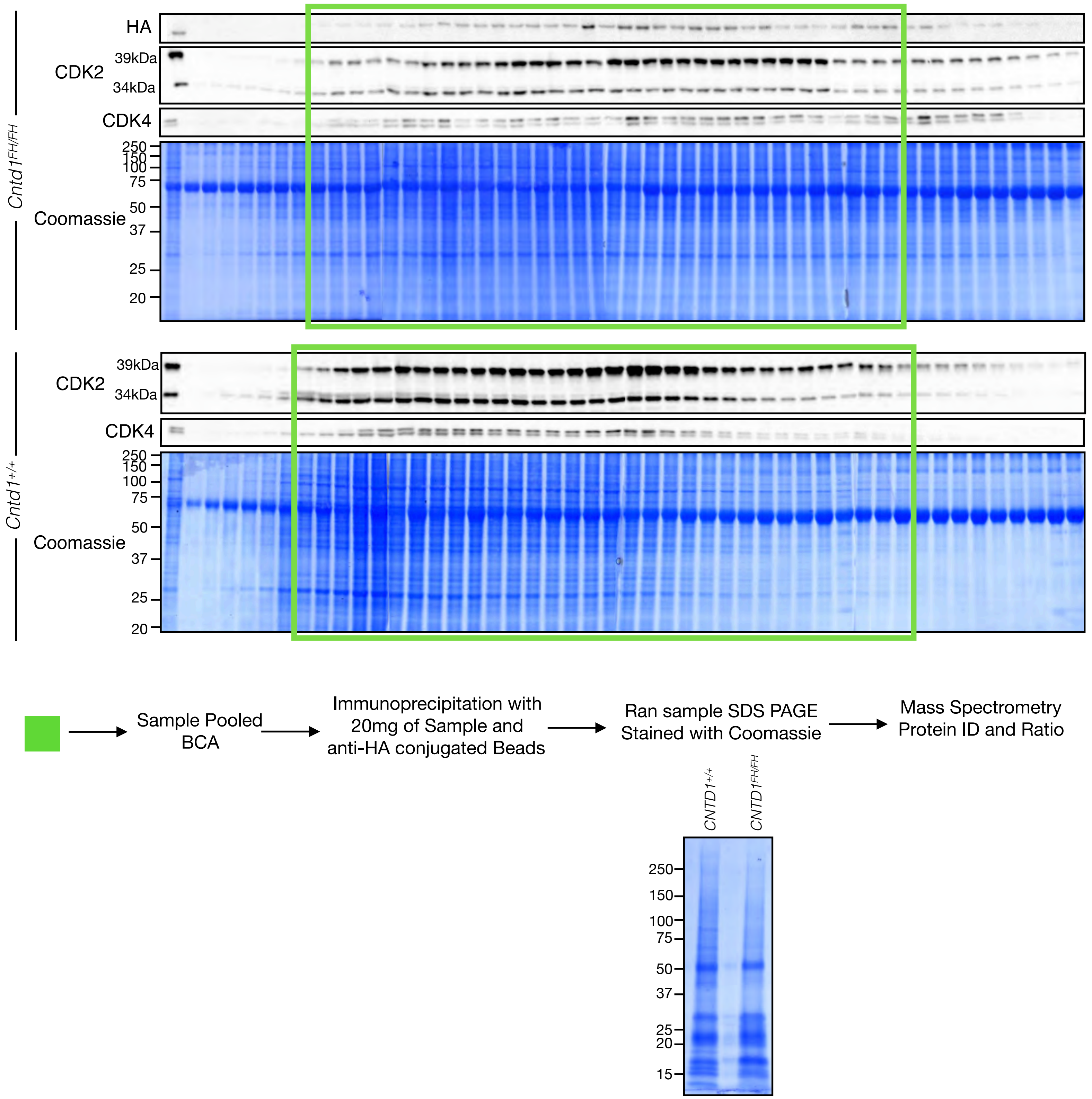
HA
CDK2
39kDa
34kDa
CDK4
Coomassie
250
150
100
75
50
37
25
20
Cntd1FH/FH
CDK2
39kDa
34kDa
CDK4
Coomassie
250
150
100
75
50
37
25
20
Cntd1+/+
Sample Pooled
BCA
Immunoprecipitation with
20mg of Sample and
anti-HA conjugated Beads
Ran sample SDS PAGE
Stained with Coomassie
Mass Spectrometry
Protein ID and Ratio
CNTD1+/+
CNTD1FH/FH
250
150
100
75
50
37
25
20
15

**a**

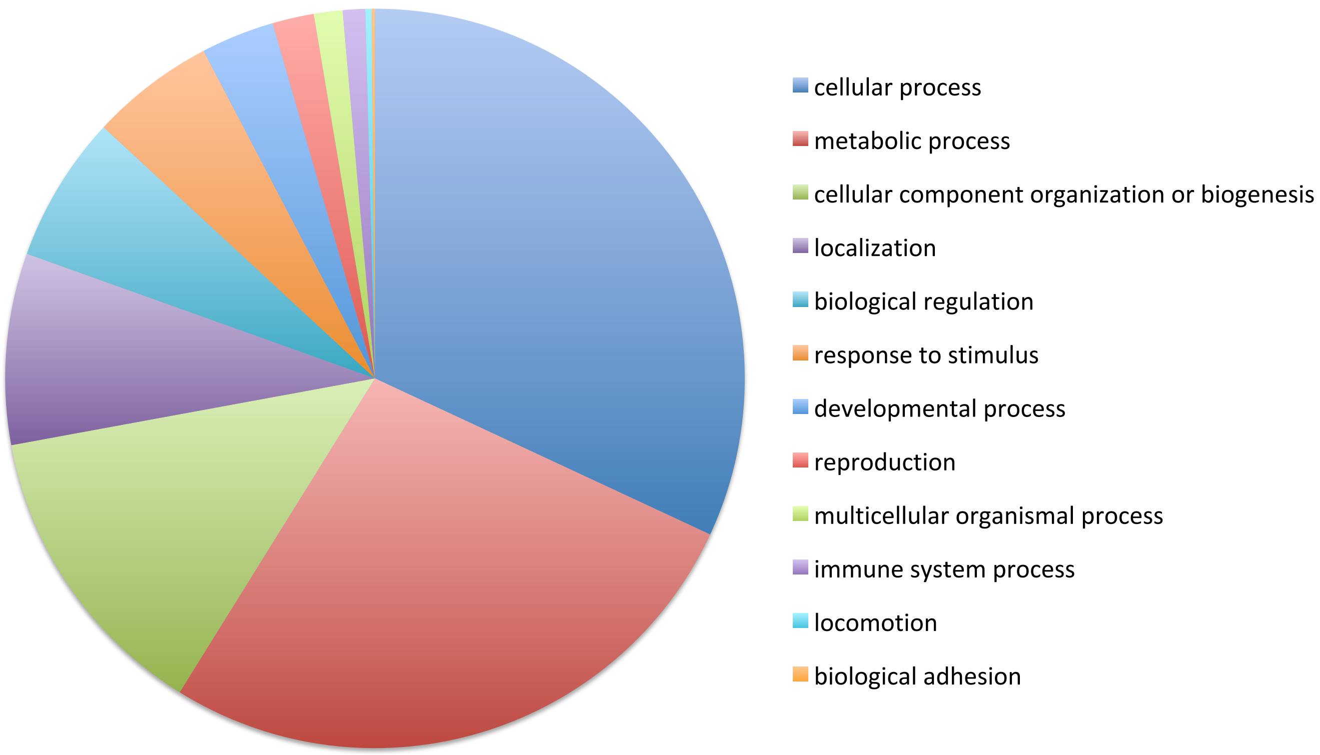


**b** CNTD1 interacting ubiquitin machinery

| E2 | E3 | De-Ubiquitin |
|---|---|---|
| ATG3 | **FBXW9** | BRCC36 |
| **CDC34** | HERC2 | USP8 |
| UBE2D3 | HERC4 | USP14 |
| | HUWE1 | USP25 |
| | NEDD4 | USP48 |
| | TM129 | VCIP135 |
| | TRIM41 | |
| | TRIM68 | |
| | UBR1 | |
| | UBR2 | |
| | UHRF1 | |
| | WWP1 | |
| | ZFP91 | |
