## Supplemental Table 2 for "Cyclin N-Terminal Domain-Containing 1 (CNTD1) coordinates meiotic crossover formation with cell cycle progression in a cyclin-independent manner"

**Supplemental Table 2: Antibody information**

| Antibody | Supplier | Catalog Number | Clone Number | Lot Number |
|---|---|---|---|---|
| **CDC34** | Proteintech | 10964-2-AP | Polyclonal | |
| **CDK1** | Millipore | MAB8878 | A17.1.1 | 3040528 |
| **CDK1-pY15** | Abcam | ab47594 | Polyclonal | GR100520-15 |
| **CDK2** | Santa Cruz | sc-163 | M2 | D2715 |
| **CDK2-pY15** | Abcam | ab76146 | EPR2233Y | GR3212813-1 |
| **CDK4** | Cell Signaling | 12790S | D9G3E | 4 |
| **Cyclin B1** | Cell Signaling | 4138S | Polyclonal | 2 |
| **FBXW9** | Thermo Fisher | PA5-70482 | Polyclonal | TH2610360 |
| **GAPDH** | Invitrogen | MA5-15738-HRP | GA1R | TJ270580 |
| **HA (Rabbit)** | Cell Signaling | 3724S | C29F4 | 9 |
| **HA (Rat)** | Roche | 11867423001 | 3F10 | 11608200 |
| **HEI10** | Abcam | ab71977 | Polyclonal | GR317016-1 |
| **MLH1** | BD Biosciences | 550838 | G168-15 | 7270524 |
| **MLH3** | Custom made previously used in (24) | | | |
| **MAD2L2** | Proteintech | 12683-1-AP | Polyclonal | |
| **Pan CDK p-Y15** | Abcam | ab133463 | EPR7875 | GR180109-14 |
| **PCNA** | Proteintech | 10205-2-AP | Polyclonal | |
| **RAD51** | Millipore | PC130-100UL | Polyclonal | 3035168 |
| **RFC3** | Proteintech | 11814-1-AP | Polyclonal | |
| **RFC4** | Proteintech | 10806-1-AP | Polyclonal | |
| **RNF212** | Novus | NBP1-83471 | Polyclonal | R34138 |
| **SYCP1** | Abcam | ab15090 | Polyclonal | GR3184119-1 |
| **SYCP3** | Custom made previously used in (24) | | | |
| **WEE1** | Novus | NBP1-33506 | Polyclonal | 42795 |
| **yH2AFX** | Millipore | 05-636 | JBW301 | 2854975 |
